# Programmable interphyla communication for *Staphylococcus aureus* by transferability of homoserine lactone sensors between Bacillales bacteria

**DOI:** 10.64898/2026.11.05.764691

**Authors:** Biprodev Sarker, Stephen N. Rondthaler, Min Zeng, Lauren B. Andrews

## Abstract

*Staphylococcus aureus* is a Gram-positive opportunistic pathogen of the order Bacillales and a frequent member of human-associated microbiomes. Despite its clinical relevance, engineering *S. aureus* and the communities it inhabits remains limited by a shortage of well-characterized genetic tools for precise control of gene expression and intercellular signaling. Biosensors for homoserine lactones (HSLs) are widely used in Gram-negative bacteria, as HSLs are small, passively diffusible signaling molecules for and their sensors are tightly regulated. Here, we show that synthetically designed HSL sensors using LuxR-family regulators are highly functional in *S. aureus*. Notably, synthetic designs previously built for the Gram-positive model *Bacillus subtilis*, a distantly related Bacillales, transferred functionally and establish repurposing the genetic toolbox as an effective strategy for importing these functions into *S. aureus*. Sensor performance correlated strongly between the two Bacillales, and activation in *S. aureus* reached 55-fold, exceeding the dynamic range of reported inducible systems. We further demonstrate tightly regulated, programmable interphyla communication between Gram-negative *Escherichia coli* and *S. aureus* via HSL signaling. These results provide a new toolset for engineering interbacterial communication and open a route to extending programmable HSL signaling to other tough-to-engineer bacteria.

## Introduction

Pathogenic *Staphylococcus aureus* can cause many diseases, often leveraging its ability to form biofilms and acquire antibiotic resistance^1,2^, yet commensal *S. aureus* strains are prevalent colonizers of human microbiomes identified in roughly one third of people^3,4^. *S. aureus* is the leading cause of hospital-associated infections^5^ and a Priority Pathogen designated by the World Health Organization^6,7^. The ability to precisely control gene regulation in *S. aureus* is crucial for understanding and developing new strategies to engineer its associated microbiomes and combat its infections^8^. Approaches from synthetic biology have proven to be apt for engineering microbial communities^9,10^. Some genetic tools have been developed for *S. aureus* (by us and other groups), including sets of graduated-strength constitutive promoters^11,12^ and inducible promoters^12–14^. However, inducible gene expression has remained moderate (up to 36-fold reported^15^), and programmable interbacterial communication to control microbiomes has not been reported for *S. aureus*. The lower genetic tractability of *S. aureus*, even compared to other Gram-positive bacteria, has hindered efforts^16^. Among the bacterial phylum Bacillota (Firmicutes), *Bacillus subtilis* is well-studied^17^ and has multiple genetic toolboxes available^18–21^. For engineering diverse bacteria, transferability of genetic parts, which is the ability of a characterized part originally built for one organism to retain its function when introduced into another organism and/or genetic context, is advantageous because it can enable standardization for scalable construction and efficient establishment of a genetic toolbox in a new organism by repurposing genetic parts across related bacterial species. In earlier work, we discovered highly functional transferability of constitutive promoters from *B. subtilis* to *S. aureus*, both members of the order Bacillales but different taxonomic families^12^. Here, we sought to test whether the genetic transferability was extensible to inducible systems for interbacterial biocommunication.

Acyl- and aryl-homoserine lactone (HSL) signals are used in natural quorum sensing to coordinate functional responses controlled by LuxR family regulators in many Gram-negative bacteria and microbiomes^22,23^. They are commonly employed in synthetic biology to engineer communication among different bacteria in order to regulate multicellular behavior and engineer microbial communities or microbial consorta^24–26^. HSLs are small diffusible molecules that bind to LuxR-family quorum sensing regulators (QSRs), enabling them to interact with specific promoters and activate transcription. Most characterized HSL sensors utilize QSRs originating from bacteria of the phylum Pseudomonadota^27^, and studies have shown that they can be engineered to function in different Gram-negative heterologous hosts^28–30^.

For interbacterial communication, HSL signaling offers several advantages over the peptide-mediated quorum-sensing systems common in Bacillales. First, HSLs are small, membrane-permeable molecules that can diffuse across the cell membrane of diverse bacteria without requiring transporters or membrane-sensing machinery necessary for peptide signals^27,31^. For example, the native Agr quorum-sensing system *S. aureus* relies on the four-gene *agrBDCA* operon, where AgrD serves as the precursor of the autoinducing peptide (AIP), AgrB participates in its processing and export, and the membrane-bound histidine kinase AgrC detects extracellular AIP and transduces the signal through the response regulator AgrA^32,33^. Moreover, because HSLs are generally heterologous to Bacillales, they offer the potential for signaling without interference with native cellular regulatory networks. A few recent studies have demonstrated that HSL sensing can be engineered in the Gram-positive bacterium, *B. subtilis*^21,34,35^. In one of these studies, we created and optimized synthetic HSL sensors for *B. subtilis*^21^ for *p*-coumaroyl-HSL (pC-HSL), 3-oxohexanoyl-HSL (3OC6-HSL), *n*-(3-hydroxytetradecanoyl)-HSL (3OHC14-HSL) and *n*-butyryl-HSL (C4-HSL)^21^. In this study, we demonstrate that synthetic HSL sensor designs are highly transferable to a distantly related Bacillales, *S. aureus*, and impart high functionality for HSL-mediated control of gene expression and tightly regulated interbacterial signaling. This work presents a simple approach of transferability of genetic systems among Bacillales bacteria to establish HSL biosensors and expand genetic tools available for *S. aureus*. We present the use of HSL sensors for bacterial interkingdom communication to a member of the Staphylococcaceae family that can be tightly controlled. Synthetic HSL sensors in Bacillales bacteria introduce the possibility of coordinating multicellular functions and interactions in engineered multispecies consortia by utilizing programmable communication between diverse bacteria utilizing HSL sensor toolboxes.

## Results

We created the genetic design for each HSL sensor containing the QSR (i.e. LuxR-family regulator) and its cognate sensor output promoter (e.g. P_Lux_). For experimental testing of the synthetic HSL sensors in *S. aureus*, we cloned the DNA constructs for four HSL sensors into *S. aureus* based on those we previously engineered for *B. subtilis* (Fig. 1a). The QSR was expressed using a strong constitutive promoter P_FtsH_ (from our genetic toolbox for *S. aureus*^12^) and original ribosome binding site (RBS) designed for *B. subtilis*. The designs included a self-cleaving ribozyme (Plmj^36^) to insulate the mRNA transcript encoding each QSR, which are transcriptional activators. To quantify sensor activity, the sensor output promoter was placed upstream of an insulated green fluorescent protein gene (*gfp*) cassette from the previously reported relative promoter unit (RPU) standard for *B. subtilis*^21^, allowing us to measure gene expression in standardized RPU units by flow cytometry with 1.0 RPU defined as the output of the strong constitutive promoter P_FtsH_ (Methods). These DNA designs for the HSL sensors come directly from our previous work in *B. subtilis*. For *B. subtilis*, DNA constructs were inserted at the *amyE* locus on the genome. For *S. aureus*, the sensor constructs were inserted into a new shuttle vector pBS9000, which is a modified version of a low-copy shuttle vector (pSR1002^12^) to here incorporate blue-white screening.

**Fig. 1:**
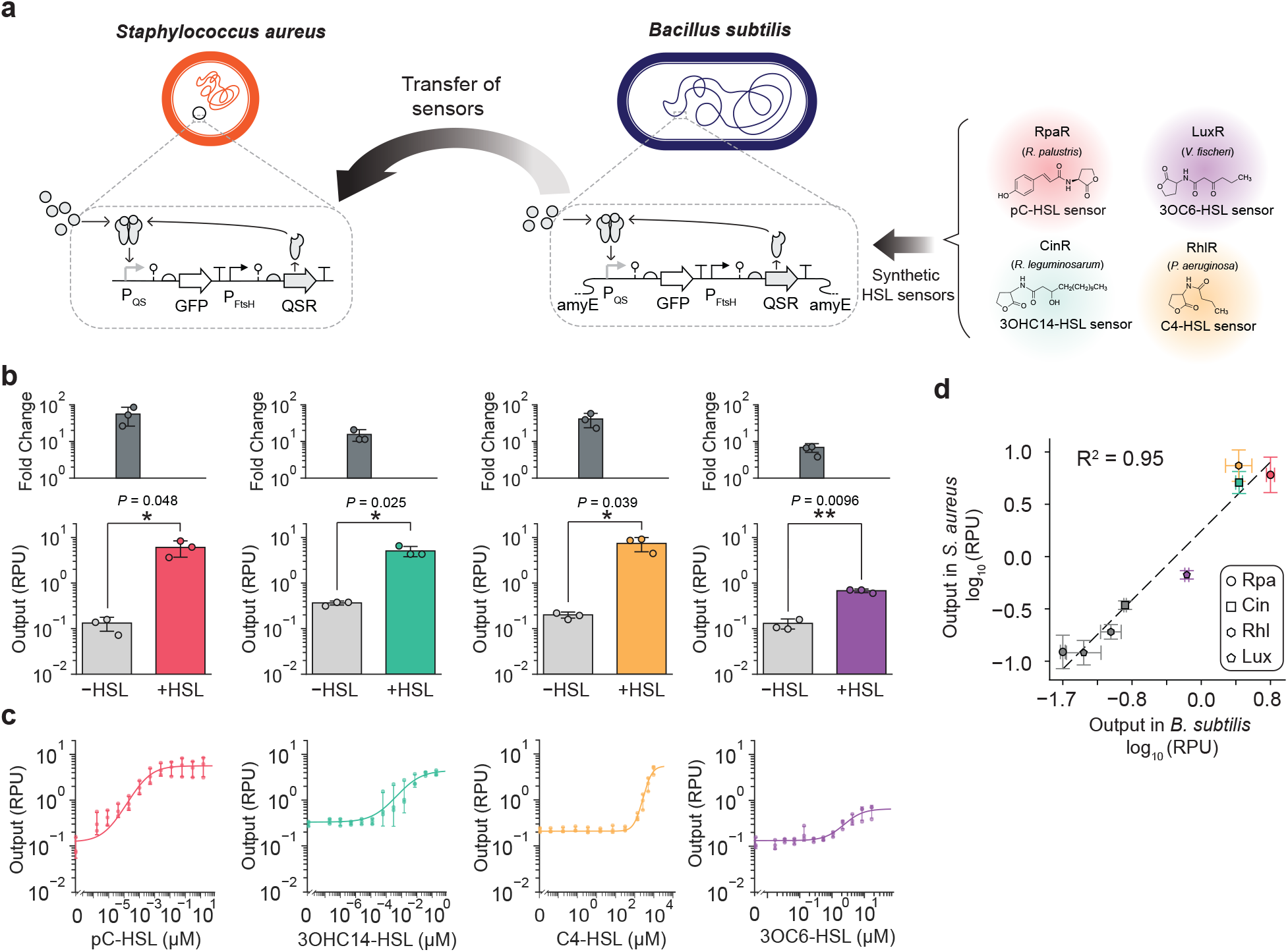
Establishing homoserine lactone (HSL) sensors for *S. aureus*. **a** Sensors were inserted on a shuttle vector for *S. aureus* (pBS9000). Each HSL sensor consists of a constitutively expressed LuxR-family regulator (RpaR, CinR, RhlR, or LuxR) that binds the cognate HSL and activates its sensor output promoter (P_QS_) to express the fluorescent reporter (GFP). HSL sensor designs built for *B. subtilis* were tested in *S. aureus*. **b** Sensor output in *S. aureus* was measured as cell fluorescence by flow cytometry (≥10,000 cells per sample). Geometric mean fluorescence was converted to standard relative promoter units (RPU) for the sensor promoter’s output (Methods). Assays were performed to measure the output without HSL (–HSL) and with 20 μM pC-HSL, 2 μM 3OHC14-HSL, 3200 μM C4-HSL, or 20 μM 3OC6-HSL (+HSL) for the sensor’s cognate ligand. Fold activation is the ratio of the measured activated output to basal output. All experiments were performed in three independent biological replicates (*n* = 3) on different days, and error bars indicate the mean ± standard deviation (S.D.). Significance was determined from a paired two-tailed Student’s t-test (*P* = 0.048 for pC-HSL, *P* = 0.025 for 3OHC14-HSL, *P* = 0.039 for C4-HSL, and *P* = 0.0096 for 3OC6-HSL sensors; * *P* < 0.05, ** *P* < 0.01). **c** Dose-response curves were assayed using intermediate HSL concentrations (*n* = 3 independent biological replicates on different days with error bars showing mean ± S.D.). The parameters for each sensor response function (line) were determined by fitting the data and are given in Supplementary Table 1. Histograms of the cell fluorescence are provided in Supplementary Fig. 1. **d** Measured activity of each sensor’s output is compared between *S. aureus* and *B. subtilis*, showing both basal output (grey) without HSL and HSL activated output (colored) at concentrations listed in panel b. The linear relationship (dotted line) determined by linear regression is plotted, and Pearson’s correlation was calculated (*n* = 3 independent biological replicates on different days, r = 0.97, *P* = 4.3×10^−5^).

To test the performance and transferability of the HSL sensors in *S. aureus*, the basal output (without HSL addition) and response to induction of the cognate HSL signal at concentrations for full activation of these sensors in *B. subtilis*^21^ (20 μM pC-HSL, 2 μM 3OHC14-HSL, 3.2 mM C4-HSL, or 20 μM 3OC6-HSL) were measured in single cells by flow cytometry. All HSL sensors exhibited significant activation in *S. aureus* (*n* = 3 independent biological replicates performed on separate days, paired two-tailed Student’s t-test, *P* = 0.048 for pC-HSL, *P* = 0.025 for 3OHC14-HSL, *P* = 0.039 for C4-HSL, and *P* = 0.0096 for 3OC6-HSL sensors). Among them, the pC-HSL sensor had the largest dynamic range, achieving 55-fold activation and low basal activity (0.12 ± 0.036 RPU (Fig. 1b). Notably, the pC-HSL sensor surpasses the fold activation of reported inducible systems for *S. aureus*, and the HSL sensors provide a new genetic tool for inducible expression.

We next characterized each HSL sensor’s dose response using intermediate concentrations of HSL that span the range from OFF to ON sensor output, in which the OFF state is without HSL, and the ON state is maximally activated (Fig. 1b). We observed smooth transition and unimodal population distributions over the range of HSL concentration (Supplementary Fig. 1). We mathematically determined each sensor’s response function, which is the input-output relationship, by fitting the data to the Hill equation that describes sensor activation (Supplementary Table 1). We also tested ligand specificity for these sensors in *S. aureus* (Supplementary Fig. 2) and found that they retain similar cross-reactivity as published earlier in *B. subtilis*^21^. We next sought to quantitatively evaluate the functional transferability of the HSLs sensors between the two bacteria. Comparing the basal and activated output for the HSL sensing systems, we observed a very strong correlation (Pearson’s r = 0.97, P = 4.3×10^−5^) for the activity between *S. aureus* and *B. subtilis* (Fig. 1c). Interestingly, this demonstrates high transferability of HSL sensing across bacterial taxonomic families (Staphylococcaceae and Bacillaceae), despite neither bacterium natively having this capability.

Altering expression of the regulator is a strategy that can tune a sensor’s response and performance^37,38^. To test tuning of the HSL sensors by increasing regulator expression, we created eight new sensor designs by substituting P_FtsH_, the constitutive promoter expressing the QSR, with two different stronger constitutive promoters previously assayed in both bacteria.^12^ The promoters P_Bla_ (native to *S. aureus*, 5.5-fold stronger than P_FtsH_ in *B. subtilis*) and P_Veg_ (native to *B. subtilis*, 12-fold stronger than P_FtsH_ in *B. subtilis*) were used. The dose response of each sensor design was characterized across a range of HSL concentrations in both *B. subtilis* (Fig. 2a) and *S. aureus* (Fig. 2b), and the corresponding sensor response functions were determined from the datasets (Supplementary Table 1). To assess whether changes in regulator expression altered the overall sensor response within each bacterial host, we compared the response functions of the three sensors containing promoter variants using a host-specific parametric functional ANOVA based on the globalized pointwise F statistic (*n* = 3 independent biological replicates performed on separate days, one-way functional ANOVA for all). In *B. subtilis*, regulator expression significantly altered the response functions of the HSL sensors containing RpaR (*P* = 9.99 × 10^−5^), RhlR (*P* = 0.0074), and LuxR (*P* = 0.00050) sensors, whereas an insignificant effect was observed for CinR (*P* = 0.520). In *S. aureus*, a significant promoter-dependent change in the overall response function was observed only for HSL sensors containing CinR (*P* = 0.0011), while response functions of sensors with RpaR (*P* = 0.280), RhlR (*P* = 0.151), and LuxR (*P* = 0.258) were not significantly different among sensor variants. Tukey-style whole-curve post-hoc comparisons corroborated the omnibus functional ANOVA results, with detailed pairwise statistical results provided in the Supplementary Data.

**Fig. 2:**
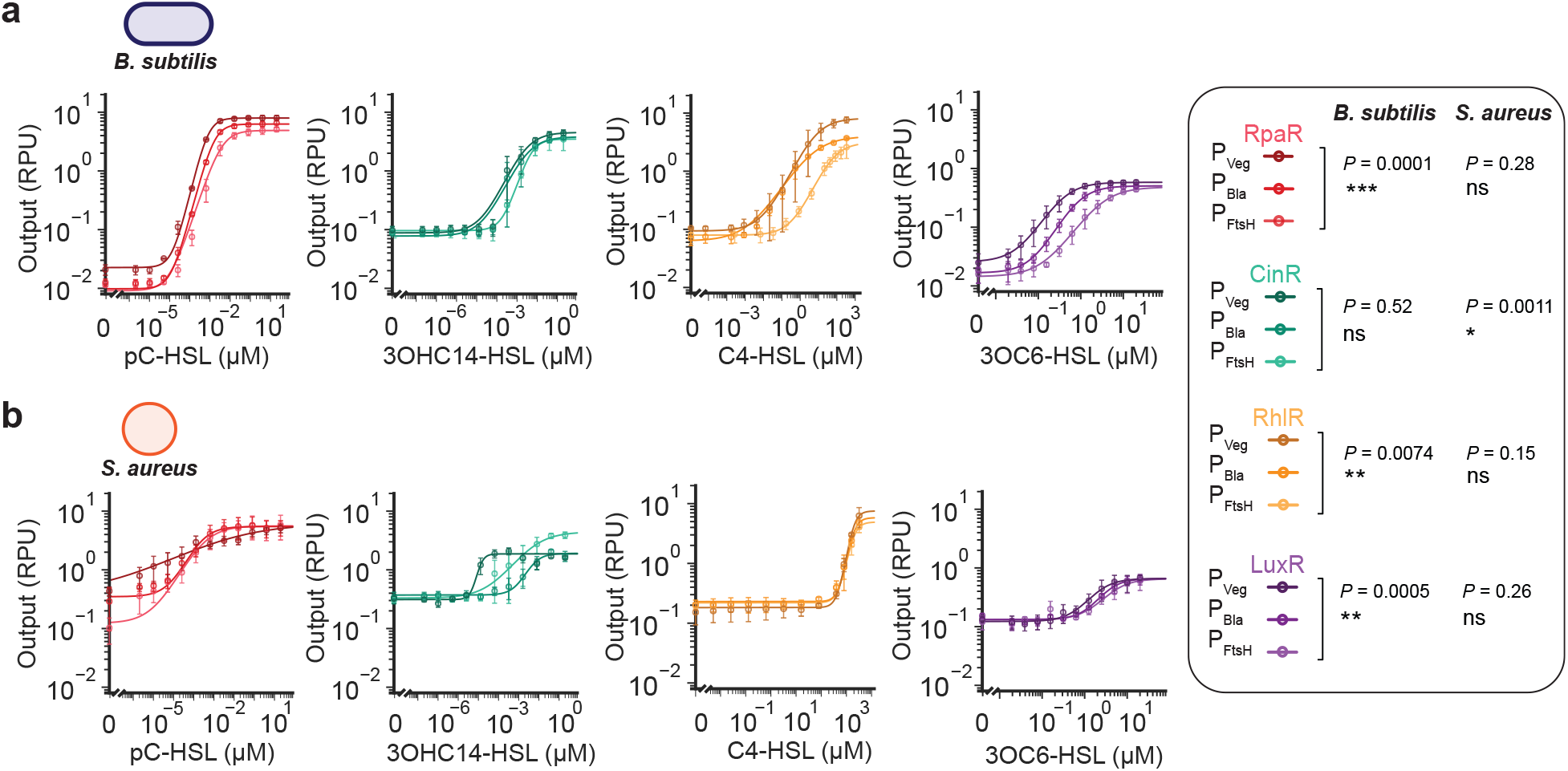
Tuning HSL sensor response using promoter variants to express quorum sensing regulators. **a** The library of twelve HSL sensor designs were generated by substituting the promoter expressing the QSR (RpaR, CinR, RhlR, LuxR) and were characterized in *B. subtilis* (*n* = 3 independent biological replicates on different days with error bars showing mean ± S.D.). **b** The same twelve HSL sensor designs were characterized in *S. aureus* to assay transferability of the genetic designs (*n* = 3 independent biological replicates on different days with errors bars showing S.D.). Sensor output as cell fluorescence was measured by flow cytometry (≥10,000 cells per sample) and converted to relative promoter units (RPU) (Methods). Statistical analysis comparing response functions for sensors containing promoter variants was performed by functional one-way ANOVA, and results are reported within the legend for *B. subtili*s and *S. aureus* (* *P* < 0.05, ** *P* < 0.01, *** *P* < 0.001, ns *P* > 0.05).

We next quantified the characteristics of the HSL biosensors based on their response functions, including: ON and OFF output levels, dynamic range (i.e. maximum fold activation), limit of detection (LOD) (defined as input yielding two-fold activation), and sensitivity (rate of change in output relative to input at the ON/OFF midpoint) for each sensor design (Fig. 3a–d). Consistent with the shifted responses observed for HSL sensors in *B. subtilis*, regulator expression significantly lowered the LOD for RpaR and LuxR (*P* = 4.6 × 10^−4^ and *P* = 0.0025, respectively) and increased sensitivity for both HSL sensors (*P* = 3.2 × 10^−4^ and *P* = 4.6 × 10^−4^, respectively) (*n* = 3 independent biological replicates on different day, one-way ANOVA analysis). Activated (ON) output also significantly increased among regulator-expression variants for RpaR and RhlR (*P* = 0.0030 and *P* = 0.0074, respectively). For example, the activated output increased from 4.88 ± 0.55 RPU (mean ± S.D., *n* = 3) to 7.96 ± 0.68 RPU while lowering the LOD from 3.67 ± 1.28 nM to 0.778 ± 0.095 nM by substituting P_FtsH_ with P_Veg_ to increase RpaR expression in the sensor design.

**Fig. 3:**
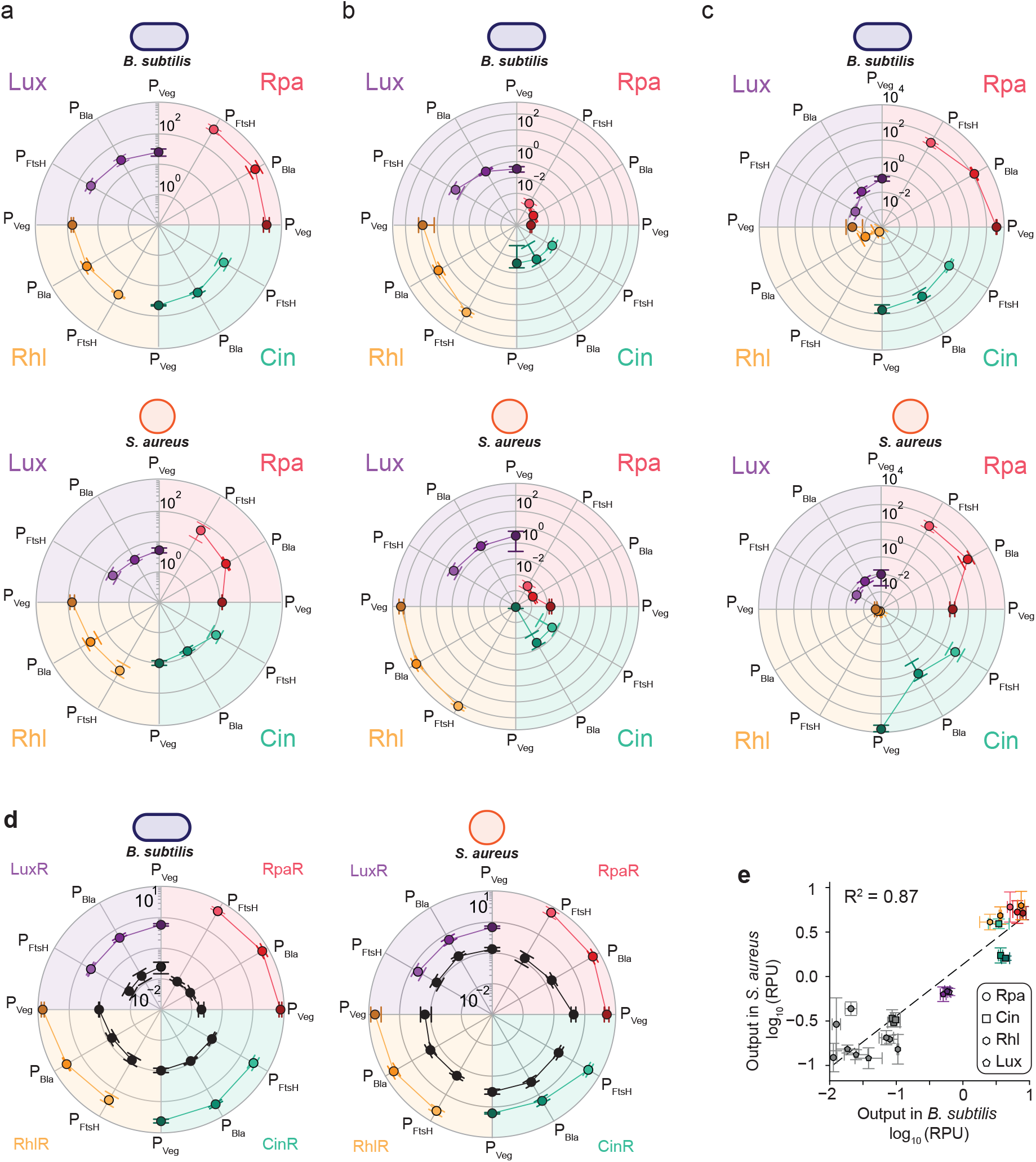
Comparing HSL sensors’ attributes between *B. subtilis* and *S. aureus*. The attributes of the HSL sensors were calculated based on the determined response function of each replicate (*n* = 3 independent biological replicates on different days). **a** Dynamic range (i.e., fold activation), **b** limit of detection (LOD), **c** sensitivity, and **d** sensor ON output (HSL induced, colored) and sensor OFF output (without HSL, black) for all 12 HSL sensor designs in *B. subtilis* and *S. aureus* is compared in radar plots. For ON state, 20 μM pC-HSL, 2 μM 3OHC14-HSL, 3200 μM C4-HSL, or 20 μM 3OC6-HSL was used for HSL induction. **e** Correlation plot compares the HSL sensors’ output (log_10_-transformed) for all twelve sensor designs between *S. aureus* and *B. subtilis*. The linear relationship (dotted line) was determined by linear regression, and Pearson’s correlation was calculated (using data from *n* = 3 independent biological replicates on different days, r = 0.93, *P* = 2.6×10^−11^). Error bars are the mean ± S.D. for the three independent experiments performed on different days in all panels.

A similarly systematic relationship between regulator expression and sensor performance was not observed in *S. aureus*. Instead, the effects were sensor specific. RpaR variants differed significantly in dynamic range (derived from *n* = 3 independent biological replicates on different day, one-way ANOVA, *P* = 0.0055), LOD (derived from *n* = 3 independent biological replicates on different day, one-way ANOVA, *P* = 4.2 × 10^−4^), and sensitivity (derived from *n* = 3 independent biological replicates on different day, one-way ANOVA, *P* = 3.5 × 10^−4^). CinR variants also differed significantly in LOD (derived from *n* = 3 independent biological replicates on different day, one-way ANOVA, *P* = 0.0039), sensitivity (derived from *n* = 3 independent biological replicates on different day, one-way ANOVA, *P* = 7.4 × 10^−4^), and activated output (derived from *n* = 3 independent biological replicates on different day, one-way ANOVA, *P* = 4.0 × 10^−4^). In contrast, changes in regulator expression produced fewer significant changes in the corresponding attributes of the RhlR and LuxR sensors. The unchanged responses for sensors containing LuxR and RhlR in *S. aureus* may be due to saturated expression in the original design. Comparing between *S. aureus* and *B. subtilis*, we observed greater uninduced basal output of the sensors (in RPU relative to P_FtsH_) in *S. aureus* than in *B. subtilis*, but not greater activated output (Fig. 3d). Nevertheless, there is a strong linear relationship for the activity of all 12 HSL sensors (4 QSRs, 3 promoters for each) between *S. aureus* and *B. subtilis* (Pearson’s r = 0.93, *P* = 2.6×10^−11^) (Fig. 3e). Therefore, while the fine-tuned sensor attributes are not retrained when transferred, the sensors retain high functionality in *S. aureus*. This demonstrates that transfer of a genetic system may quickly yield a functional starting point, yet optimization, if necessary, should be conducted in the bacterial host.

HSL sensing is natively used to mediate communication and interactions between bacteria in a microbiome. Therefore, we next sought to investigate whether we could program *S. aureus* to receive and respond to HSL signals produced by another bacterium using the developed sensors. For this purpose, we utilized *E. coli* strains that produce 3OHC14-HSL, 3OC6-HSL, and C4-HSL via isopropyl β-D-thiogalactopyranoside (IPTG) inducible expression of the corresponding HSL synthase (CinI, LuxI, or RhlI, respectively), using the P_Tac_ promoter to regulate HSL biosynthesis. Interbacterial signaling was assayed using the *S. aureus* strains for HSL sensing when grown in media of the *E. coli* sender cultures or wild-type (WT) *E. coli* as negative control. We observed that all three cell-cell communication systems showed activated output in *S. aureus* with induction of HSL signal biosynthesis, with up to 9.2-fold activation for the Cin system (Fig. 4c), whereas no sensor activation was observed when WT *E. coli* was used. This demonstrates programmable HSL-mediated interbacterial communication to *S. aureus* and across bacterial kingdoms (Pseudomonadati to Bacillati) can be achieved using these HSL sensors for *S. aureus*.

**Fig. 4:**
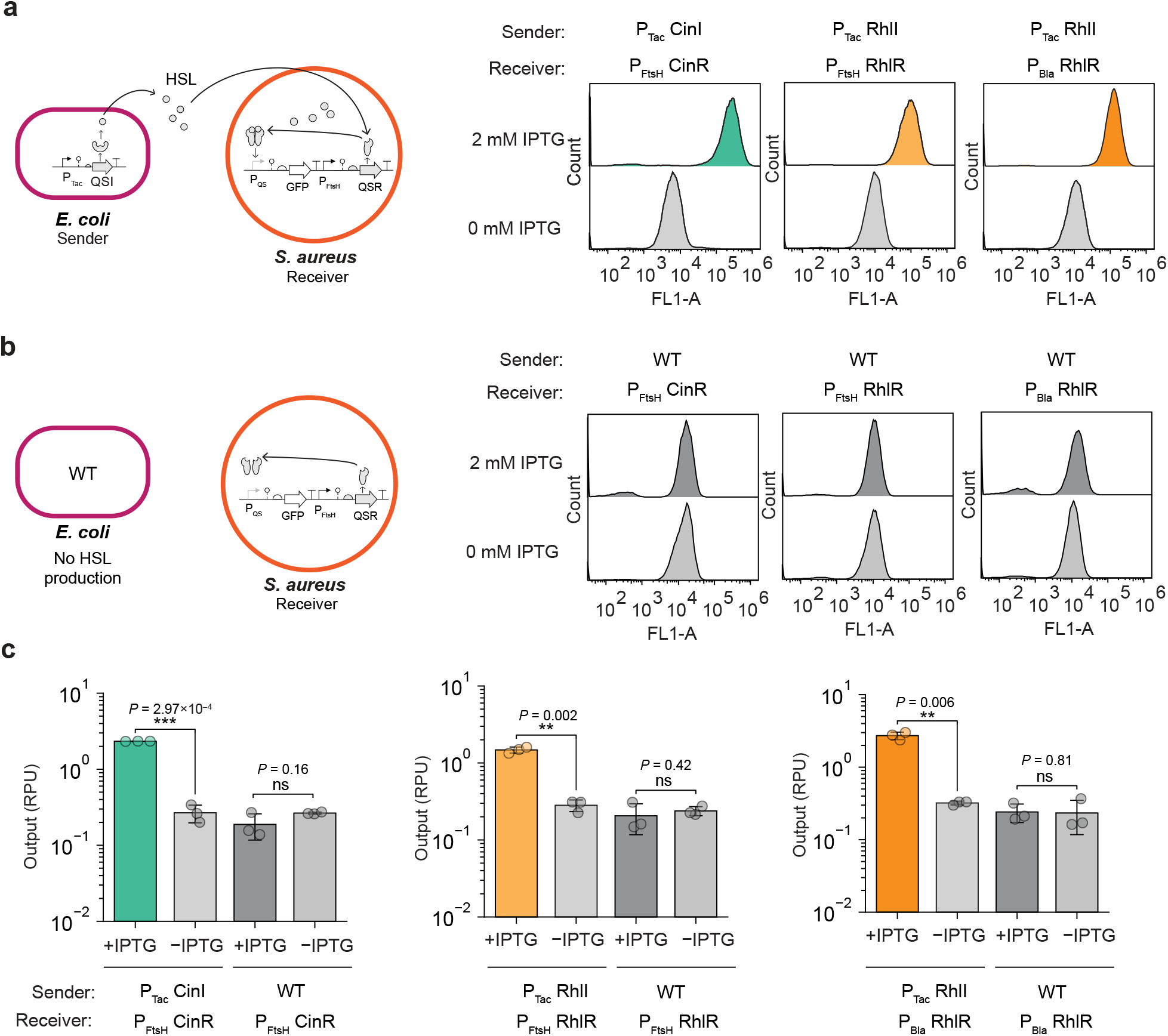
Programmable interbacterial signaling using *S. aureus* containing synthetic HSL sensors. **a** HSL sender strains of *E. coli* contained IPTG-inducible expression of the corresponding HSL synthase (QSI = CinI, RhlI, or LuxI), and *S. aureus* receiver strains contained a corresponding synthetic HSL sensor. *S. aureus* was cultured with supernatants of the sender strains grown with 2 mM IPTG induction (+ IPTG) or without IPTG (– IPTG) addition. Output of the HSL sensor in *S. aureus* was assayed by flow cytometry in three identical experiments performed on separate days (≥10,000 cells per sample), and representative histograms of HSL detection in *S. aureus* are shown. All histograms for all sensors with CinR, RhlR, and LuxR are provided in Supplementary Figure 3. **b** Control experiments were performed identically using the wildtype (WT) *E. coli* in place of the engineered *E. coli* sender strains, and representative histograms are shown. **c** The measured sensor output was converted to relative promoter units (RPU) (Methods). Markers show the values from each of the 3 independent biological replicates performed on different days in addition to mean ± S.D. (*n* = 3). Statistical significance of sensor activation was calculated using paired two-tailed Student’s t-tests comparing the conditions with and without IPTG induction of HSL synthase for each sender-receiver strain pair (** *P* < 0.01, *** *P* < 0.001, ns *P* > 0.05) (*n* = 3 independent biological replicates on separate days).

## Discussion

This study establishes the capability for HSL-mediated communication to be engineered in Bacillales bacteria that do not naturally harbor these systems. The work identifies transferability of synthetic genetic designs for HSL sensing between distantly related Bacillales bacteria (here using *B. subtilis* and *S. aureus*) as an effective approach to impart new functions for programmable bacterial communication. This finding of shared transferability among the Bacillales bacteria suggests that the strategy may be effective for other bacterial relatives, including non-model strains. Here, a library of biosensors to detect four diverse acyl-(C4-C12) and aryl-homoserine lactones was established for both

S. *aureus* and *B. subtilis* and utilized design variants to alter QSR regular expression to tune the sensor’s properties. Given that the synthetic HSL sensors were transferable using four different QSRs (RpaR, RhlR, CinR, LuxR), this suggests that the approach likely could be applied more broadly for HSL sensors utilizing various LuxR-family regulators. The synthetic HSL sensors expands and surpasses reported inducible systems for S. *aureus* with up to 55-fold activation (compared to 28-fold^12^ and 36-fold activation reported^15^) and presents further improved HSL sensors for *B. subtilis* beyond the initial reported designs^11^. This work also demonstrates bacterial interkingdom (from Pseudomonadati to Bacillati) programmable communication to a member of the Staphylococcaceae family that can be tightly controlled using different diffusible HSLs and synthetic HSL sensors. Whereas HSL-mediated programmable interbacterial communication has been limited to Gram-negative bacteria, here we show that the synthetic HSL sensors are useful in Bacillales to be able to engineer communication and interactions between Bacillales in bacterial consortia comprised of diverse bacteria and not limited to communication between Gram-negative bacteria.

HSL-mediated signaling offers many advantages for programming communication and interspecies functions within bacterial communities and consortia comprised of widely varied bacteria. Of note, HSL signaling requires relatively few components and is genetically compact. Typically, a single QSR regulator is needed for an HSL sensor, facilitating its construction and tuning. In contrast, peptide-based communication commonly used by Gram-positive bacteria requires peptide processing and export, membrane-bound sensing, and two-component signal transduction, such as the native Agr system in *S. aureus*. A recent study demonstrated the feasibility of transferring the Agr system to *E. coli*, achieving an approximately 4-fold dynamic range following optimization of multiple circuit components^39^. By using membrane-diffusible signals, HSL-mediated communication circumvents the requirement for many of these components.

The results of this study promote HSL signaling as an attractive platform for engineering communication among various bacteria by utilizing the high transferability and large dynamic range of various HSL sensors in heterologous hosts. We envision that programmable HSL-mediated communication with Bacillales bacteria could be applied for a wide range of applications, such as engineered communication among Staphylococci in the skin microbiome or programmable communication among therapeutic bacterial consortia used to control delivery of payloads. Looking ahead, further study is warranted to determine the extensibility of the observed genetic transferability to other Firmicutes and to establish the limits for engineering diverse microbiomes. The approach and toolboxes presented in this study provide a foundation for controlling HSL-mediated communication in Bacillales and diverse bacteria that has the potential to be broadly applied in synthetic biology for programmable interbacterial interactions.

## Methods

### Strains and plasmids

*Staphylococcus aureus* ATCC 12600 was acquired from BEI Resources (Manassas, VA USA) and used for all testing in *S. aureus. Bacillus subtilis* 168 was acquired from the Bacillus Genetic Stock Center (Columbus, OH USA) and used for all testing in *B. subtilis*. Cloning was performed in the *Escherichia coli* NEB 10-beta strain from New England Biolabs (NEB) (Ipswich, MA USA) and subsequently transformed into *E. coli* IM08B^40^ (from BEI Resources) prior to transformation to *S. aureus* to ensure compatible methylation. All plasmids are listed in Supplementary Table 2, and plasmid maps are in Supplementary Fig. 4. All DNA primers and oligos were purchased from Integrated DNA Technologies (Coralville, IA USA). All genetic part DNA sequences are listed in Supplementary Table 3 and Supplementary Data. Oligos are listed in Supplementary Table 4. Plasmids, including the HSL sensors, are available from Addgene (Watertown, MA USA).

### Media and reagents

For cloning and propagation, all strains were cultured in Miller’s LB broth (Fisher Bioreagents) purchased from Thermo Fisher Scientific (Waltham, MA USA) and on LB agar plates. *S. aureus* strains were grown in SSM9PR minimal media^41^ for assaying sensor activities. All chemicals were purchased from MilliporeSigma (Burlington, MA USA) unless otherwise noted. SSM9PR contained 1X M9 salts (6.78 g/L Na_2_HPO_4_, 1.0 g/L NH_4_Cl, 3.0 g/L KH_2_PO_4_, and 0.5 g/L NaCl, 2 mM MgSO_4_, 0.1 mM CaCl_2_, 1% w/v D-glucose, 1% w/v casamino acids (Acros), 1 mM thiamine hydrochloride (Fisher Bioreagents), and 0.05 mM nicotinamide (Acros). Media used for assaying sensors in *B. subtilis* contained 1X M9 salts supplemented with 0.34 g/L thiamine hydrochloride, 0.2% w/v casamino acids, 2 mM MgSO_4_, 0.4% w/v D-glucose, 0.1 mM CaCl_2_, and 20 µg/ml L-tryptophan (for tryptophan auxotrophy).

*B. subtilis* 168 and *S. aureus* ATCC 12600 were made competent using SpC and SpII media. SpC media contained T-base (15 mM (NH_4_)_2_SO_4_, 80 mM K_2_HPO_4_, 44 mM KH_2_PO_4_, and 3.4 mM Na_3_C_6_H_5_O_7_) and was supplemented with 1 mM MgSO_4_, 0.5% w/v D-glucose, 0.2% w/v yeast extract, 0.025% w/v casamino acids, and 20 µg/ml L-tryptophan. SpII media contained T-base supplemented with 3.5 mM MgSO_4_, 0.5% w/v D-glucose, 0.1% w/v yeast extract, and 0.01% w/v casamino acids. For transformations,

*E. coli* and *B. subtilis* strains were recovered in SOC media (20 g/L tryptone, 5 g/L yeast extract, 10 mM MgCl_2_, 20 mM MgSO_4_, 10 mM NaCl, and 2.5 mM KCl). *S. aureus* strains were recovered in B2 recovery media containing 10 g/L casamino acids, 25 g/L yeast extract, 1 g/L K_2_HPO_4_, 5 g/L glucose, and 25 g/L NaCl. Blue-white screening used agar plates overlaid with 80 µL 20 mg/mL X-gal (GoldBio) and 40 µL 100 mM isopropyl β-D-thiogalactopyranoside (IPTG, GoldBio) purchased from GoldBio (St. Louis, MO USA).

Antibiotic was added to media for plasmid maintenance, and all antibiotics were purchased from GoldBio. For *E. coli* strains, the following antibiotic concentrations were used for plasmid maintenance: 100 µg/mL carbenicillin, 50 µg/mL kanamycin, and 400 µg/mL erythromycin. For *S. aureus*, 10 µg/mL erythromycin was used for plasmid maintenance. For engineered *B. subtilis* strains, 5 µg/mL chloramphenicol was used to select for transformed cells. Homoserine lactones were purchased from MilliporeSigma for *n*-butyryl-HSL (C4-HSL) (Cat. 09945), 3-oxohexanoyl-HSL (3OC6-HSL) (Cat. K3007), *n*-(3-hydroxytetradecanoyl)-HSL (3OHC14-HSL) (Cat. 51481), and *p*-coumaroyl-HSL (pC-HSL) (Cat. 07077). HSL stock solutions were dissolved in pure DMSO (Fisher Bioreagents). Chloramphenicol and erythromycin were dissolved in absolute ethanol (Fisher BioReagents), and other chemicals were aqueous stock solutions.

### Plasmid DNA construction

The plasmid pBS9000 was used as the backbone for the sensor designs for *S. aureus*. pBS9000 was constructed by Type IIS (Golden Gate) DNA assembly using PCR products from the pSR1002^12^ vector and *lacZα* gene for blue-white screening. All sensors for *S. aureus* were assembled using hierarchical Type IIS DNA assembly into this vector as described in our previous study^21^. Briefly, transcriptional unit constructs were assembled using BsaI-HFv2 (NEB) for the sensor output promoter expressing *gfp* and for a constitutive promoter expressing the QSR gene in separate assembly reactions. The two transcription units for each HSL sensor were inserted into pBS9000 in a subsequent DNA assembly reaction using BbsI (NEB). To construct sensor design variants for *B. subtilis*, the original constitutive promoter (P_FtsH_ ) on the integration plasmid for each sensor^21^ was removed using inverse PCR. Primers appended BbsI recognition sequences for the insertion of P_Bla_ or P_Veg_ as annealed ssDNA oligos with appropriate linker sequences. All Type IIS DNA assembly reactions contained: 10 fmol of DNA insert if plasmid or 100 fmol of annealed oligos or purified PCR products, 5 fmol of destination vector, 5 U Type IIS restriction enzyme (BbsI, SapI or BsaI-HFv2), 250 U of T4 DNA ligase (NEB), 1X T4 ligase buffer, and nuclease-free water for a 5 µL total reaction volume. DNA assembly reactions were incubated in a thermal cycler from Bio-Rad(Hercules, CA USA) using the protocol: alternating between 37 °C and 16 °C for 2.5 min each for 36 cycles, followed by 50 °C for 30 min for digestion of unassembled vector, and then enzyme inactivation for 20 min at 65 °C (for BbsI and SapI) or 80 °C (for BsaI).

Each DNA assembly reaction was transformed into *E. coli* NEB 10-beta cells. For transformation, 1.5 µL of assembly reaction was mixed with 5 µL chemically competent cells. Transformations were plated on LB agar for blue-white screening with erythromycin for *S. aureus* sensors or with kanamycin for *B. subtilis* sensors. One white colony from each transformation was cultured for 16 h at 37 °C and 250 rpm in an Eppendorf Innova 44 (Enfield, CT USA) shaking incubator in LB with antibiotic, and plasmid DNA was purified using a Qiagen miniprep kit (Germantown, MD USA). All assembled DNA constructs were verified using Sanger sequencing (Azenta) and/or whole plasmid DNA sequencing (Plasmidsaurus). Plasmids designed for *S. aureus* were subsequently transformed into chemically competent *E. coli* IM08B to achieve proper methylation prior to transformation into *S. aureus*.

### *B. subtilis* 168 competent cell preparation and transformation

To prepare competent cells of *B. subtilis*, single colony from a freshly streaked plate was used to inoculate 5 mL LB medium in a 14 mL snap-cap culture tube and grown at 37 °C with shaking at 250 rpm for 16 h. The overnight culture was then diluted into 12.5 mL pre-warmed SpC medium to an initial OD_600_ of 0.01 in a 125 mL Erlenmeyer flask and incubated at 37 °C and 250 rpm with OD_600_ measurements taken every 30 min. Upon reaching the onset of stationary phase, the culture was incubated for an additional 2 h, followed by a 10-fold dilution into 25 mL pre-warmed SpII medium in a 250 mL Erlenmeyer flask and further incubation at 37 °C and 250 rpm for 1.5 h. Cells were harvested at room temperature by centrifugation at 10,000 G, concentrated 20-fold in the remaining SpII medium, and stored at −70 °C in 10% v/v glycerol for later use.

For transformation of *B. subtilis*, approximately 1 µg purified integration plasmid was added to 100 µL competent cells in a culture tube and incubated at 37 °C with shaking at 250 rpm for 1 h, followed by the addition of 500 µL SOC and a further 2 h incubation under the same conditions to allow recovery. Cells were then concentrated 5-fold by centrifugation at 10,000 G and plated onto LB agar containing chloramphenicol. Resulting colonies were streak-purified twice serially by overnight growth on chloramphenicol plates. Genome integration was confirmed for a subset of constructs by long-read DNA sequencing (Plasmidsaurus) of PCR amplicons of the *amyE* locus amplified from purified genomic DNA.

### *S. aureus* electrocompetent cell preparation and transformation

To prepare electrocompetent cells of *S. aureus*, a freshly streaked colony was used to inoculate 5 mL of LB medium in a culture tube and incubated at 37 °C and 250 rpm for 16 h overnight. Next, the culture was diluted into 12.5 mL prewarmed SpC media to an OD_600_ 0.01 in a 125 mL flask. The OD_600_ was measured every 30 min during incubation at 250 rpm and 37 °C, and the culture was incubated until 2 h after reaching stationary phase. The culture was diluted 10-fold into 25 mL warm SpII media in a 250 mL flask and incubated at 250 rpm and 37 °C for 1.5 h. Next, the cells were placed in an ice–water bath for 10 min, then transferred to 50 mL conical tubes and centrifuged at 10,000 G for 10 min at 4 °C using a fixed-angle rotor. The supernatant was decanted, and the pellet was gently resuspended in 50 mL ice-cold sterile Milli-Q water while maintained on ice, followed by an additional 3–5 min chilling period before centrifugation. This washing procedure was repeated twice, with cells resuspended sequentially in 25 mL ice-cold sterile MilliQ water and then 12.5 mL sterile 10% v/v glycerol. After the final spin, the pellet was gently resuspended in 500 µL ice-cold sterile 10% v/v glycerol and portioned into 100 µL aliquots for immediate electroporation or stored at −70 °C for future use.

For transformation of *S. aureus*, a 100 µL aliquot of electrocompetent cells was thawed on ice and mixed with approximately 1 µg plasmid DNA purified from the *E. coli* IM08B strain. The cell–DNA suspension was incubated on ice for 30 min, then transferred to a pre-chilled 1 mm electroporation cuvette (Fisher Scientific) and electroporated at 2.1 kV with a pulse length of 1.1 ms using a Bio-Rad MicroPulser. Immediately following electroporation, 500 µL of pre-warmed B2 medium was added directly to the cuvette, mixed thoroughly, and transferred to a 1.5 mL microcentrifuge tube for recovery at 37 °C with shaking at 250 rpm for 1 h. Recovered cells were plated onto LB agar containing antibiotic and incubated overnight at 37 °C.

### Sensor characterization assays

All sensors were assayed using flow cytometry. *S. aureus* strains containing a sensor characterization plasmid were streaked on LB agar plates. One fresh colony for each strain was picked to inoculate 200 µL SSM9PR in a 96-well U-bottom microtiter plate and covered with a breathable seal (Excel Aeraseal) before incubating in a plate shaking incubator ELMI DTS-4 (Newbury Park, CA USA) at 37 °C and 1000 rpm for 16 h. After overnight incubation, cultures in the plate were diluted 1600-fold (5 µL culture into 195 µL fresh SSM9PR for two serial dilutions) and incubated 3 h in the 96-well plate. After this growth, cells were diluted 40-fold (5 µL of culture into 195 µL media) into SSM9PR with addition of HSL as needed and incubated in a 96-well plate for 21 h. Next, the *S. aureus* cells were fixed using 4% v/v paraformaldehyde in phosphate buffered saline (PBS at pH 7.4) incubated at room temperature for 0.5 h^12^. Fixed cells were diluted in PBS (∼1600-fold) prior to flow cytometry.

Sensors in *B. subtilis* were assayed using the identical conditions as for *S. aureus*, except a couple differences. *B. subtilis* was cultured in M9 media with L-tryptophan instead of SSM9PR medium. *B. subtilis* cells were not fixed prior to flow cytometry. Instead, *B. subtilis* cultures were diluted (∼1600-fold) into PBS (pH 7.4) with 2 mg/mL kanamycin to stall cell growth and incubated at room temperature for 0.5 h prior to flow cytometry.

### Interbacterial signaling assay

For interbacterial HSL signaling to *S. aureus*, assays were performed similarly to sensor characterization. To prevent competitive growth, bacteria were cultured separately. For the HSL sender, *E. coli* 10-beta (NEB) strains containing IPTG-inducible expression of the corresponding HSL synthase (sender) were streaked on LB agar with kanamycin and incubated overnight. For control experiment, WT *E. coli* 10-beta (NEB) were streaked on LB agar without any antibiotic. One *E. coli* colony for each strain was picked to inoculate 200 µL M9 media (identical to M9 for *B. subtilis* except without tryptophan) with kanamycin for engineered *E. coli* or without any antibiotic for WT *E. coli* in a 96-well U-bottom plate covered with a breathable seal and incubated at 37 °C and 1000 rpm for 16 h. After overnight growth, the culture was diluted 1200-fold by inoculating 5 µL of overnight culture into 6 mL M9 with kanamycin for engineered strains or without any antibiotic and, if needed, 2 mM IPTG to induce HSL synthase expression and incubated for 24 h in a 14 mL culture tube. The supernatant was harvested by centrifugation (10,000 G for 15 min) to pellet the cells. *S. aureus* receiver cells were prepared identical to the sensor characterization, except 100 µL of the supernatants of the *E. coli* strains were mixed with 95 µL of fresh SSM9PR with erythromycin and added to 5 µL *S. aureus* culture in the final dilution plate. After 21 h incubation, flow cytometry was performed as described above.

### Flow cytometry analysis

Cell fluorescence was measured using a BD Accuri C6 (Franklin Lakes, NJ USA) flow cytometer equipped with a 488 nm blue laser and 533/30 nm emission filter for GFP. Samples were collected at the medium flow rate (66 µL/min) using the FL1-A channel. For each sample, at least 10,000 gated cell events (Supplementary Fig. 5) were collected. Data were analyzed using the FlowJo software (v10). A cell gate was created for each bacterium to distinguish cells from electronic noise or debris using the wildtype strain for each. The geometric mean fluorescence of cells was determined and converted to relative promoter units (RPU) for the sensor output promoter activity. This allows for the output promoter strength to be determined relative to a standard promoter and compared across instruments and labs. The conversion to RPU sensor output was calculated using the equation:

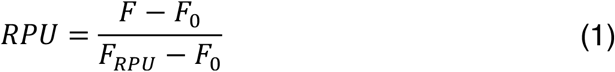

where the quantities are the fluorescence of the sample cells (*F*), the autofluorescence of the wildtype cells (*F*_*0*_), and fluorescence of cells harboring the RPU standard construct (*F*_*RPU*_). The RPU standard construct for *B. subtilis* expresses *gfp* via the constitutive promoter P_FtsH_ from the *amyE* locus using integration plasmid pMZ6000 (Supplementary Fig. 4). For *S. aureus*, the RPU standard plasmid (pBS9100) is the shuttle vector containing the identical transcriptional unit for constitutive GFP expression with P_FtsH_ (Supplementary Fig. 4). Accordingly, 1.0 RPU is equal to the promoter strength of P_FtsH_ in the genetic context used for measuring the sensor output. The RPU standard construct has been previously published for *B. subtilis* 168^21^. The wildtype cells and RPU standard construct for each bacterium were assayed in parallel for flow cytometry performed on each day.

### Data and statistical analysis

To determine the response function for each HSL sensor, the measured sensor output data was fit to the Hill equation for activation as a function of the input HSL concentration. The fitting was performed using SciPy^42^ in a custom Python script using the least squares method. The Hill equation use is:

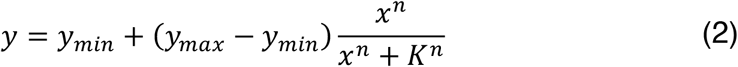

where *y* is the sensor output (in RPU), and *x* is the input concentration of HSL (in *µ*M). The Hill parameters are: *y*_*min*_ basal output (in RPU), *y*_*max*_ maximum activated output (in RPU), *K* Hill dissociation constant (in *µ*M), and *n* Hill coefficient (unitless).

To compare the sensor properties in both bacteria, the response function and its parameters for each experimental replicate were determined (Supplementary Data). The activated output (*y*_*max*_) and basal sensor output (*y*_*min*_) were used directly (in RPU). The dynamic range was calculated as the ratio of the activated and basal sensor output (*y*_*max*_/*y*_*min*_). Limit of detection (in μM cognate HSL) was determined as *x* the input HSL concentration (in *µ*M) resulting in two-fold greater output than the basal output (2*y*_*min*_). The sensitivity of the sensor (in RPU/μM cognate HSL) is the slope of the sensor response function (*dy/dx*) evaluated at the mid-threshold (*x* where y is the average of *y*_*max*_ and *y*_*min*_).

### Statistics and Reproducibility

All experiments were performed in three independent replicates performed on separate days. For statistical analysis of sensor activation, paired two-tailed Student’s t-tests were performed between sensor outputs in RPU of the uninduced and induced output (*n* = 3). The linear regression of log10 transformed values were performed using a custom Python script using the LinearRegression() function of sklearn.linear_model^43^ package. Pearson’s correlation to compare the sensor outputs in *S. aureus* and *B. subtilis* was performed using the SciPy package (v1.17.0) in Python and scipy.stats.pearsonr() function. The functional ANOVA and Tukey-style whole-curve post-hoc analyses were implemented in custom Python scripts using NumPy. One-way ANOVA was performed using scipy.stats.f_oneway() in SciPy. Conventional Tukey HSD tests for sensor attributes were performed in Python’s statsmodels library using the function statsmodels.stats.multicomp.pairwise_tukeyhsd(). Data and statistical results are provided in the Supplementary Data.

### Data availability

All data generated and analyzed during this study are included in this published article and its supplementary information files. The source data for all figures is provided in the Supplementary Data file.

## Acknowledgements

This study was funded by the U.S. National Science Foundation (Grant No. MCB-2211039, Grant No. DMR-1904901, Grant No. CBET-1943695) and the Marvin and Eva Schlanger faculty fellowship to LBA. SNR was supported by the NSF Graduate Research Fellowship Program (Award No. DGE-1451512).

## Contributions

B.S., S.R., and L.B.A. conceived the project and designed the experiments. B.S., S.R., and M.Z. performed the experiments. B.S. and L.B.A. analyzed the data. L.B.A. supervised the project. B.S. and L.B.A. wrote the draft. All authors reviewed and approved the manuscript.

## Ethics declarations

The authors declare no competing interests.

