## Supplementary Information for "Programmable interphyla communication for *Staphylococcus aureus* by transferability of homoserine lactone sensors between Bacillales bacteria"

**Supplementary Figures:**
Supplementary Figure 1. Cell fluorescence histograms for HSL sensor response curves in *S. aureus*
Supplementary Figure 2. Signal crosstalk characterization of HSL sensors in *S. aureus*
Supplementary Figure 3. Interbacterial signaling to *S. aureus* for sensor library
Supplementary Figure 4. Plasmid maps
Supplementary Figure 5. Gating strategy
Supplementary Figure 6. Histograms of *S. aureus* WT and RPU standard

**Supplementary Tables:**
Supplementary Table 1. Parameters for response functions of HSL sensors
Supplementary Table 2. Plasmids used in this work
Supplementary Table 3. Genetic part sequences used in this work
Supplementary Table 4. DNA oligos used in this work

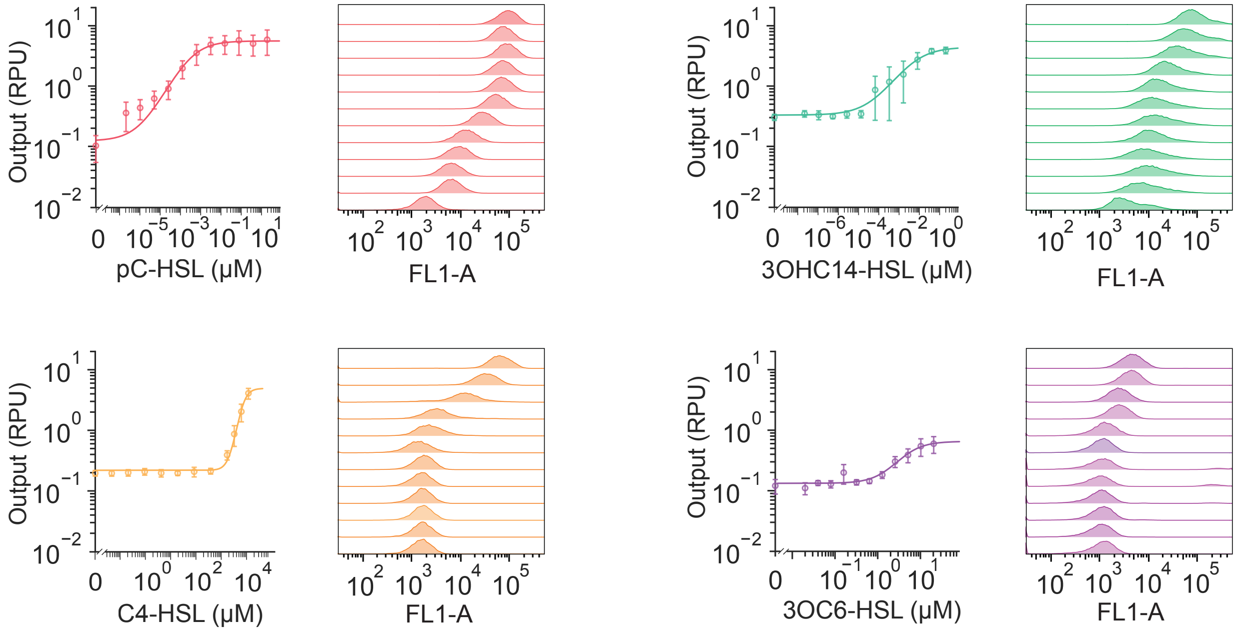


**Supplementary Fig. 1: Cell fluorescence histograms for HSL sensor response curves in *S. aureus*.** Dose response curves for each sensor with $P_{FtsH}$ expressing a QSR are shown. One representative histogram is shown for each point on the dose response curve. Sensor output as cell fluorescence was measured by flow cytometry (≥10,000 cells per sample). The geometric mean fluorescence in arbitrary units was converted to relative promoter units (RPU) for the sensor promoter's output relative to a standard (Methods). The markers show the average of three independent experiments performed on different days, and error bars are the mean ± standard deviation (SD) ($n$ = 3 for biologically independent experiments on separate days).

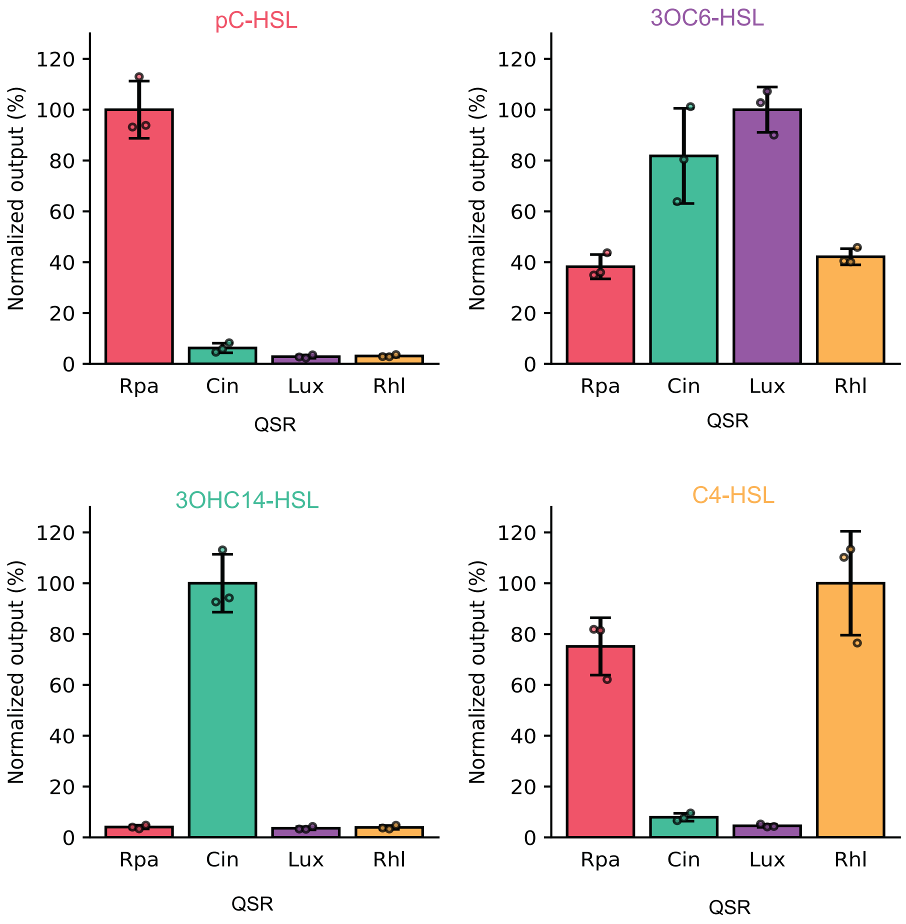


**Supplementary Fig. 2: Signal crosstalk characterization of HSL sensors in *S. aureus*.** QSRs are expressed by $P_{FtsH}$ for each sensor. Each sensor was induced by cognate and three noncognate HSLs. The concentration of HSLs were chosen based on the full activation of each sensor: 20 µM for pC-HSL, and 3OC6-HSL; 2 µM for 3OHC14-HSL; and 3200 µM C4-HSL. The cell fluorescence was measured by flow cytometry (≥ 10,000 cells per sample). The geometric mean of cell fluorescence in arbitrary units was converted to standard relative expression units (RPU). Bars are the average of normalized RPU in percent output with respect to the cognate HSL sensor output of three independent experiments performed on different days. Error bars are the mean ± SD (*n* = 3 for biologically independent experiments on separate days).

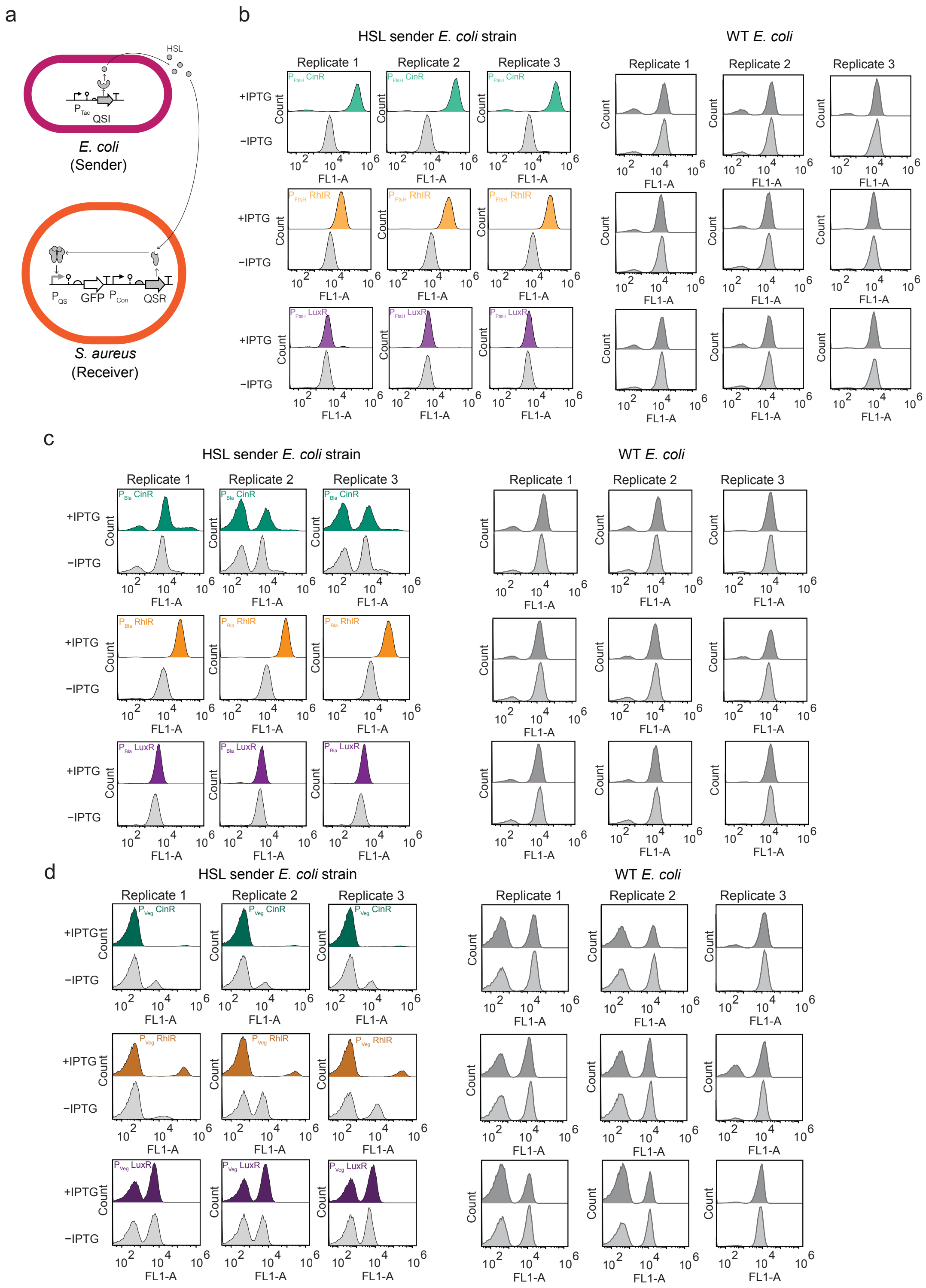


**Supplementary Fig. 3: Interbacterial signaling to *S. aureus* for sensor library. (a)** Genetic schematic for HSL sensing in *S. aureus* of HSL produced by *E. coli*. Histograms are the *S. aureus* output from three biologically independent replicates of the assay performed on different days. *S. aureus* HSL sensors with QSR (CinR, LuxR, or RhlR) expressed by **(b)** $P_{FtsH}$ **(c)** $P_{Bla}$ **(d)** $P_{Veg}$ were assayed with supernatant from the appropriate *E. coli* HSL-producing strain with 2 mM IPTG induction (colors) or without induction of the HSL synthase (CinI, LuxI, or RhlI), or S. aureus grown with WT *E. coli* supernatants without and with 2 mM IPTG induction.

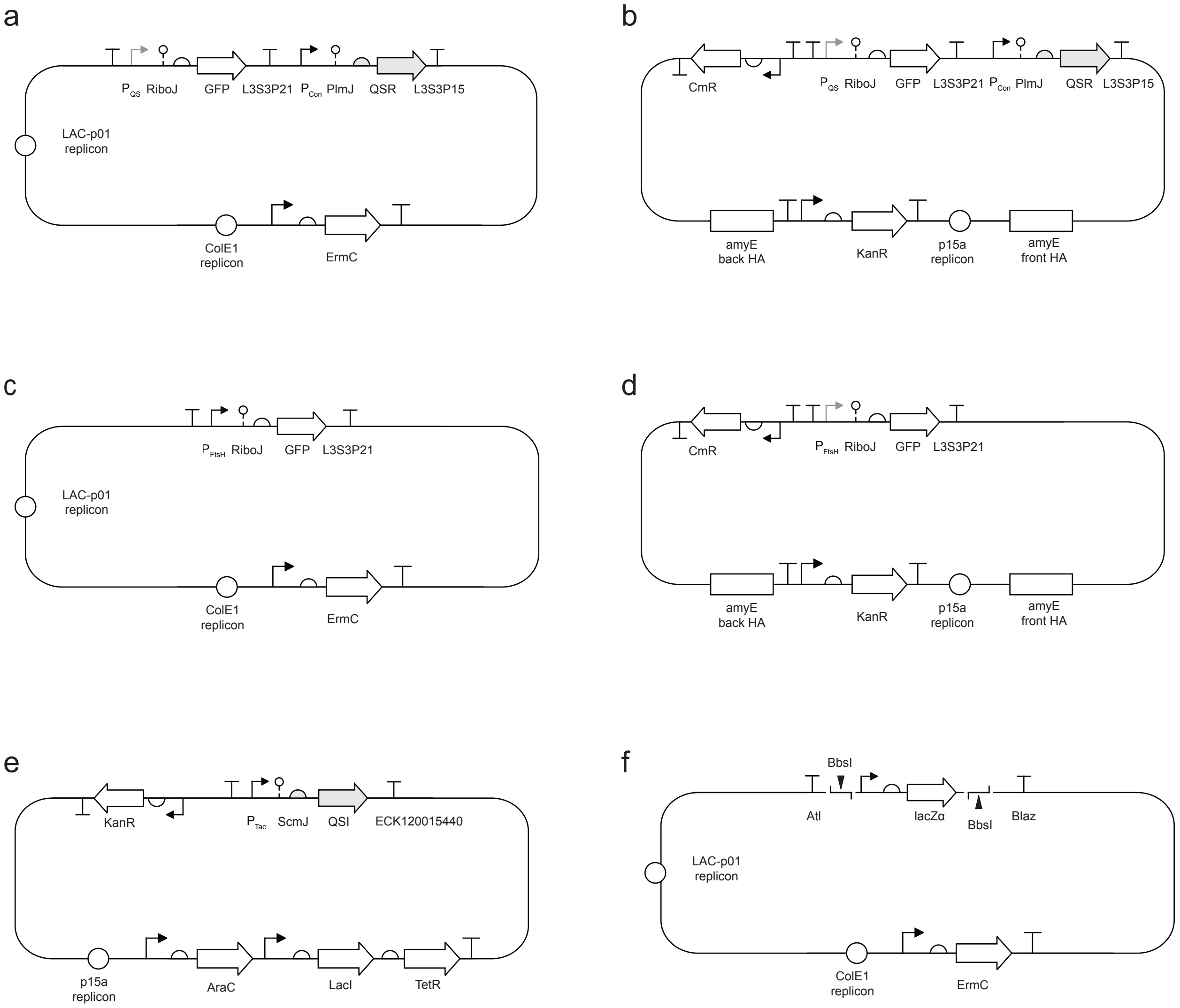


**Supplementary Fig. 4: Plasmid maps. (a)** For *S. aureus*, genetic schematic for plasmids for sensor characterization. **(b)** For *B. subtilis*, genetic schematic for integration plasmids containing the sensor characterization construct flanked by homology arms for *amyE* genomic integration of the sensor constructs. For all HSL sensors, the QSRs were RpaR, LuxR, RhlR, or CinR with the corresponding $P_{QS}$ ($P_{Rpa}$, $P_{Lux}$, $P_{Rhl}$, or $P_{Cin}$, respectively). $P_{Con}$ was $P_{FtsH}$, $P_{Bla}$, or $P_{Veg}$ for each corresponding sensor variant. **(c)** *S. aureus* RPU standard plasmid (pBS9100). **(d)** Integration plasmid for *B. subtilis* RPU standard strain. **(e)** HSL bioproduction plasmids for *E. coli* expressing the QSI (CinI, RhlI, or LuxI). **(f)** pBS9000 *E. coli*-*S. aureus* shuttle vector for constructing sensor designs for *S. aureus.*

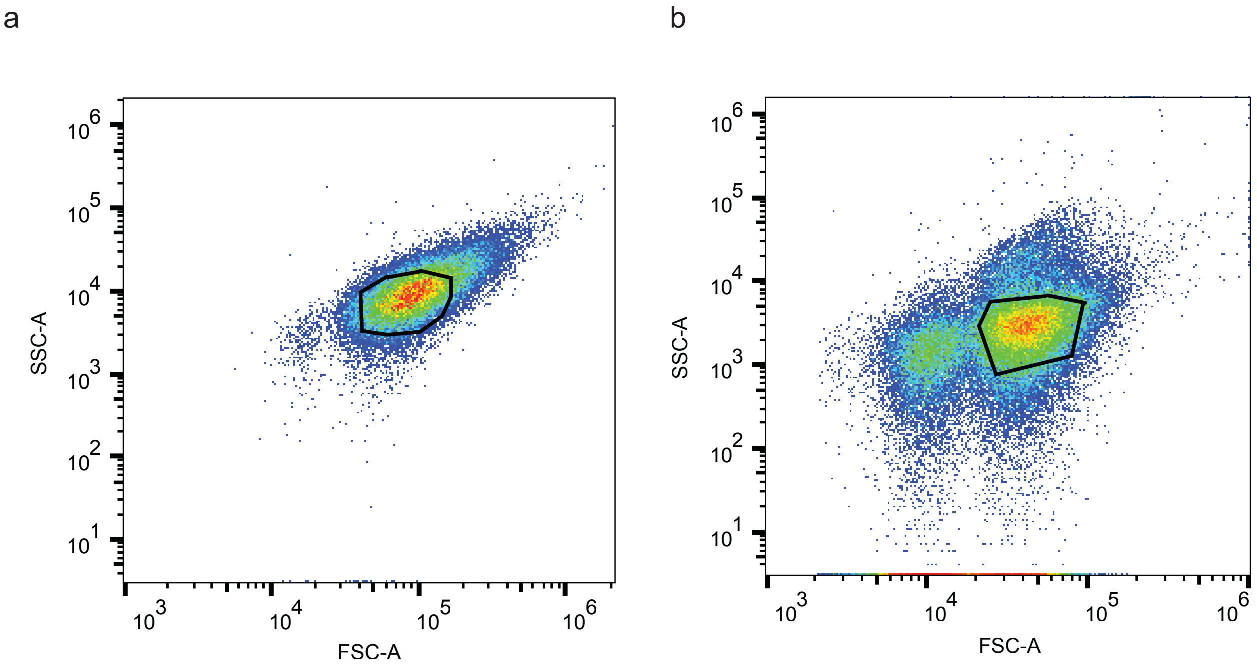


**Supplementary Fig. 5: Gating strategy.** Flow cytometry samples were gated around the visible focus in the SSC-A vs FSC-A scatter plot in FlowJo. The examples shown here are for **(a)** the *S. aureus* RPU standard strain, and **(b)** the *B. subtilis* RPU standard strain.

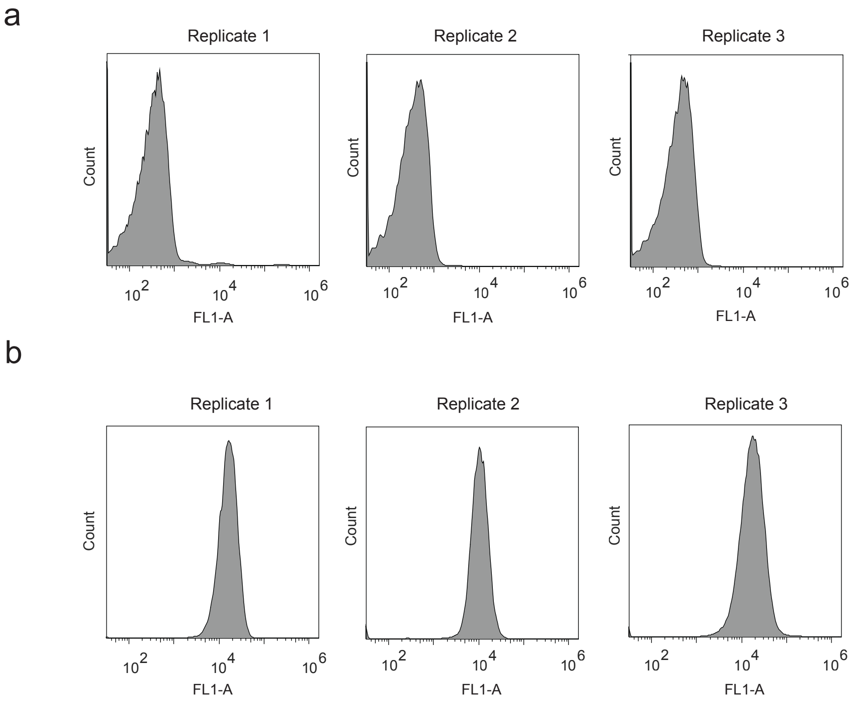


**Supplementary Fig. 6: Histograms of *S. aureus* WT and RPU standard.** Flow cytometry histograms of **(a)** wildtype (WT) *S. aureus* and **(b)** the *S. aureus* RPU standard cells in the FL1-A fluorescence channel. Plots from three biologically independent experiments ($n$ = 3) performed on separate days are shown (≥ 10,000 cells per sample).

**Supplementary Table 1. Parameters for response functions of HSL sensors**

| QSR | QSR promoter | Bacterium | K (μM) | n | $y_{min}$ (RPU) | $y_{max}$ (RPU) |
|---|---|---|---|---|---|---|
| RpaR | $P_{FtsH}$ | *B. subtilis* | 0.0036 | 1.07 | 0.01 | 4.9 |
| RpaR | $P_{Bla}$ | *B. subtilis* | 0.0016 | 1.33 | 0.01 | 6.29 |
| RpaR | $P_{Veg}$ | *B. subtilis* | 0.0008 | 1.46 | 0.02 | 7.96 |
| CinR | $P_{FtsH}$ | *B. subtilis* | 0.0415 | 1.26 | 0.1 | 3.49 |
| CinR | $P_{Bla}$ | *B. subtilis* | 0.031 | 0.79 | 0.08 | 3.81 |
| CinR | $P_{Veg}$ | *B. subtilis* | 0.026 | 0.81 | 0.09 | 4.53 |
| LuxR | $P_{FtsH}$ | *B. subtilis* | 2.91 | 1.19 | 0.01 | 0.48 |
| LuxR | $P_{Bla}$ | *B. subtilis* | 0.82 | 1.47 | 0.02 | 0.51 |
| LuxR | $P_{Veg}$ | *B. subtilis* | 0.37 | 1.38 | 0.03 | 0.58 |
| RhlR | $P_{FtsH}$ | *B. subtilis* | 509.45 | 0.76 | 0.08 | 3.06 |
| RhlR | $P_{Bla}$ | *B. subtilis* | 44.4 | 0.52 | 0.06 | 3.92 |
| RhlR | $P_{Veg}$ | *B. subtilis* | 64.35 | 0.69 | 0.09 | 7.95 |
| RpaR | $P_{FtsH}$ | *S. aureus* | 0.0031 | 0.68 | 0.12 | 5.6 |
| RpaR | $P_{Bla}$ | *S. aureus* | 0.0021 | 0.86 | 0.35 | 5.47 |
| RpaR | $P_{Veg}$ | *S. aureus* | 0.039 | 0.23 | 0.39 | 6.19 |
| CinR | $P_{FtsH}$ | *S. aureus* | 0.043 | 0.6 | 0.33 | 4.43 |
| CinR | $P_{Bla}$ | *S. aureus* | 0.042 | 1.22 | 0.37 | 1.91 |
| CinR | $P_{Veg}$ | *S. aureus* | 0.0001 | 2.59 | 0.31 | 1.87 |
| LuxR | $P_{FtsH}$ | *S. aureus* | 4.58 | 1.52 | 0.13 | 0.65 |
| LuxR | $P_{Bla}$ | *S. aureus* | 3.31 | 1.76 | 0.13 | 0.66 |
| LuxR | $P_{Veg}$ | *S. aureus* | 2.05 | 1.6 | 0.12 | 0.66 |
| RhlR | $P_{FtsH}$ | *S. aureus* | 1833.34 | 2.53 | 0.22 | 4.91 |
| RhlR | $P_{Bla}$ | *S. aureus* | 1845.46 | 2.68 | 0.23 | 5.81 |
| RhlR | $P_{Veg}$ | *S. aureus* | 1855.21 | 2.77 | 0.18 | 7.53 |

**Supplementary Table 2. Plasmids used in this work**

| Plasmid | Description | Reference |
|---|---|---|
| pSR1002 | Engineered *E. coli-S. aureus* shuttle vector with erythromycin antibiotic resistance, ColE1 origin of replication for *E. coli*, and LAC-p01 replicon for *S. aureus*. The vector was built from pKK30[1]. | [2] |
| pBS9000 | *E. coli-S. aureus* shuttle vector identical to pSR1002, except with *lacZα* between BbsI restriction sites for blue-white screening of assembled constructs in *E. coli* cloning strain. | This study |
| pBS9001 | pC-HSL sensor characterization construct for *S. aureus* on shuttle vector pBS9000. $P_{Rpa}$ expresses GFP. RpaR expressed by $P_{FtsH}$ and RpaR RBS. | This study |
| pBS9002 | 3OHC14-HSL sensor characterization construct for *S. aureus* on shuttle vector pBS9000. $P_{Cin}$ expresses GFP. CinR expressed by $P_{FtsH}$ and CinR RBS. | This study |
| pBS9003 | 3OC6-HSL sensor characterization construct for *S. aureus* on shuttle vector pBS9000. $P_{Lux}$ expresses GFP. LuxR expressed by $P_{FtsH}$ and LuxR RBS. | This study |
| pBS9004 | C4-HSL sensor characterization construct for *S. aureus* on shuttle vector pBS9000. $P_{Rhl}$ expresses GFP. RhlR expressed by $P_{FtsH}$ and LuxR RBS. | This study |
| pBS9006 | pC-HSL sensor characterization construct for *S. aureus* on shuttle vector pBS9000. $P_{Rpa}$ expresses GFP. RpaR expressed by $P_{Bla}$ and RpaR RBS. | This study |
| pBS9007 | 3OHC14-HSL sensor characterization construct for *S. aureus* on shuttle vector pBS9000. $P_{Cin}$ expresses GFP. CinR expressed by $P_{Bla}$ and CinR RBS. | This study |
| pBS9008 | 3OC6-HSL sensor characterization construct for *S. aureus* on shuttle vector pBS9000. $P_{Lux}$ expresses GFP. LuxR expressed by $P_{Bla}$ and LuxR RBS. | This study |
| pBS9009 | C4-HSL sensor characterization construct for *S. aureus* on shuttle vector pBS9000. $P_{Rhl}$ expresses GFP. RhlR expressed by $P_{Bla}$ and LuxR RBS. | This study |
| pBS9010 | pC-HSL sensor characterization construct for *S. aureus* on shuttle vector pBS9000. $P_{Rpa}$ expresses GFP. RpaR expressed by $P_{Veg}$ and RpaR RBS. | This study |

| | | |
|---|---|---|
| pBS9011 | 3OHC14-HSL sensor characterization construct for *S. aureus* on shuttle vector pBS9000. $P_{Cin}$ expresses GFP. CinR expressed by $P_{Veg}$ and CinR RBS. | This study |
| pBS9012 | 3OC6-HSL sensor characterization construct for *S. aureus* on shuttle vector pBS9000. $P_{Lux}$ expresses GFP. LuxR expressed by $P_{Veg}$ and LuxR RBS. | This study |
| pBS9013 | C4-HSL sensor characterization construct for *S. aureus* on shuttle vector pBS9000. $P_{Rhl}$ expresses GFP. RhlR expressed by $P_{Veg}$ and LuxR RBS. | This study |
| pBS9100 | RPU standard plasmid for *S. aureus*. GFP is constitutively expressed by $P_{FtsH}$ and GFP RBS on pBS9000 backbone; ribozyme riboJ used for insulating GFP mRNA. | This study |
| pNH446 | RPU standard integration plasmid for integration at the *amyE* locus for *B. subtilis*. GFP is constitutively expressed by $P_{FtsH}$ and GFP RBS; ribozyme RiboJ used for insulating GFP mRNA. | [3] |
| pMZ1001 | pC-HSL sensor construdtion for integration at *amyE* locus of *B. subtilis*. $P_{Rpa}$ expresses GFP. RpaR expressed by $P_{FtsH}$ and RpaR RBS. | [3] |
| pMZ2001 | 3OC6-HSL sensor construdtion for integration at *amyE* locus of *B. subtilis*. $P_{Lux}$ expresses GFP. LuxR expressed by $P_{FtsH}$ and LuxR RBS. | [3] |
| pMZ3001 | C4-HSL sensor construdtion for integration at *amyE* locus of *B. subtilis*. $P_{Rhl}$ expresses GFP. RhlR expressed by $P_{FtsH}$ and RhlR RBS. | [3] |
| pMZ4001 | 3OHC14-HSL sensor construdtion for integration at *amyE* locus of *B. subtilis*. $P_{Cin}$ expresses GFP. RpaR expressed by $P_{FtsH}$ and CinR RBS. | [3] |
| pBS9101 | pC-HSL sensor construdtion for integration at *amyE* locus of *B. subtilis*. $P_{Rpa}$ expresses GFP. RpaR expressed by $P_{Bla}$ and RpaR RBS. | This study |
| pBS9102 | 3OHC14-HSL sensor construdtion for integration at *amyE* locus of *B. subtilis*. $P_{Cin}$ expresses GFP. RpaR expressed by $P_{Bla}$ and CinR RBS. | This study |

| | | |
|---|---|---|
| pBS9103 | 3OC6-HSL sensor construdtion for integration at *amyE* locus of *B. subtilis*. $P_{Lux}$ expresses GFP. LuxR expressed by $P_{Bla}$ and LuxR RBS. | This study |
| pBS9104 | C4-HSL sensor construdtion for integration at *amyE* locus of *B. subtilis*. $P_{Rhl}$ expresses GFP. RhlR expressed by $P_{Bla}$ and RhlR RBS. | This study |
| pBS9105 | pC-HSL sensor construdtion for integration at *amyE* locus of *B. subtilis*. $P_{Rpa}$ expresses GFP. RpaR expressed by $P_{Veg}$ and RpaR RBS. | This study |
| pBS9106 | 3OHC14-HSL sensor construdtion for integration at *amyE* locus of *B. subtilis*. $P_{Cin}$ expresses GFP. RpaR expressed by $P_{Veg}$ and CinR RBS. | This study |
| pBS9107 | 3OC6-HSL sensor construdtion for integration at *amyE* locus of *B. subtilis*. $P_{Lux}$ expresses GFP. LuxR expressed by $P_{Veg}$ and LuxR RBS. | This study |
| pBS9108 | C4-HSL sensor construdtion for integration at *amyE* locus of *B. subtilis*. $P_{Rhl}$ expresses GFP. RhlR expressed by $P_{Veg}$ and RhlR RBS. | This study |
| pMZ306 | 3OC6-HSL bioproduction using LuxI signal biosynthesis module ($P_{Tac}$-LuxI) in *E. coli* | This study |
| pMZ307 | C4-HSL bioproduction using RhlI signal biosynthesis module ($P_{Tac}$-RhlI) in *E. coli* | This study |
| pMZ309 | 3OHC14-HSL bioproduction using LuxI signal biosynthesis module ($P_{Tac}$-CinI) in *E. coli* | This study |

**Supplementary Table 3. Genetic part sequences used in this work**

| Part Name | Type | DNA sequence | Reference |
|---|---|---|---|
| $P_{FtsH}$ | Promoter | ATGGTTATTGTTTGTATTGGAATGATTTTCTATGGTACTATTGAACATAGTTGTGA | 4 |
| $P_{Bla}$ | Promoter | TTGACACCGATATTACAATTGTAATATTATTGATTT | 2 |
| $P_{Veg}$ | Promoter | TTGATATAATTTAAATTTTATTTGACAAAAATGGGCTCGTGTTGTACAATAAATGT | 2 |
| $P_{Rpa}$ | Promoter | GCACCTGTCCGATCGGACAGTATTACGCAAGAAAATGGTTTGTTATAGTAAATGTAGT | 3 |
| $P_{CinD}$ | Promoter | CATGCTCTGATCCCCCTCATCTGGGGGGCCTATCTGAGGGAAATTACGCAAGAAAATGGTTATGTTATAGTAAATGTAGT | 3 |
| $P_{Lux}$ | Promoter | ACCTGTAGGATCGTACAGGTATTACGCAAGAAAATGGTTTGTTATAGTAAATGTAGT | 3 |
| $P_{Rhl}$ | Promoter | TCCTGTGAAATCTGGCAGTTACCGTTAGCTTTCGAATTGGCTATAATAAATGTAGT | 3 |
| RiboJ | Insulator | AGCTGTCACCGGATGTGCTTTCCGGTCTGATGAGTCCGTGAGGACGAAACAGCCTCTACAAATAATTTTGTTTAA | 5 |
| PlmJ | Insulator | AGTCATAAGTCTGGGCTAAGCCCACTGATGAGTCGCTGAAATGCGACGAAACTTATGACCTCTACAAATAATTTTGTTTAA | 5 |
| GFP RBS | RBS | TACTAGAGGAAGCTTACATAAGGAGGAACTACT | 3 |
| RpaR RBS | RBS | TACGACTGCGAGGTCATAAGAAAGGAGGTTACGTT | 3 |
| LuxR RBS | RBS | TACGCCATATAAAAAGGAGACACTACCTCA | 3 |
| RhlR RBS | RBS | TACGTTAGGGAGTAAGTAAAGGAGGTAAAATATT | 3 |
| CinR RBS | RBS | TACGTCCGTACATAGCAGAAGGGGGTAAATA | 3 |
| CinI RBS | RBS | AAAAAAAAAGCATTGATAG | This study |
| LuxI RBS | RBS | GCTATACGTTCATAAGGAGGTATTTTC | This study |
| RhlI RBS | RBS | GGCATAGGAGGACACTCTTA | This study |
| *rpaR* | CDS | ATGATCGTCGGCGAAGATCAGCTTTGGGGACGGCGTGCGCTGGAATTCGTCGATTCCGTCGAACGGCTCGAGGCGCCGGCGCTGATCAGCCGGTTCGAATCGCTGATCGCGAGCTGCGGATTTACCGCCTAC | 6 |

| | | | |
|---|---|---|---|
| | | ATCATGGCCGGCCTGCCGTCGCGCAA<br>TGCCGGACTACCGGAGCTGACGCTGG<br>CCAATGGCTGGCCGCGAGACTGGTTC<br>GATCTGTATGTCAGCGAAAACTTCAGC<br>GCGGTCGATCCGGTGCCGCGCCACGG<br>CGCTACCACGGTTCATCCTTTCGTATG<br>GTCCGATGCACCCTACGACCGCGACC<br>GTGATCCGGCCGCCCACCGGGTCATG<br>ACCCGGGCGGCGGAATTCGGACTGGT<br>CGAGGGTTACTGCATTCCGCTGCACTA<br>CGACGACGGTAGCGCCGCGATCAGCA<br>TGGCCGGCAAGGATCCGGACCTCAGC<br>CCGGCCGCGCGCGGCGCGATGCAGC<br>TGGTCAGCATCTACGCGCATAGTCGCC<br>TGCGCGCACTCAGCCGGCCAAAGCCG<br>ATCCGGCGCAACCGGCTCACGCCGCG<br>CGAGTGCGAGATCCTGCAATGGGCAG<br>CGCAGGGCAAGACCGCCTGGGAAATC<br>TCGGTAATCCTCTGCATCACCGAACGC<br>ACGGTGAAATTCCATCTGATCGAAGCC<br>GCCCGCAAGCTCGACGCCGCCAACCG<br>CACCGCGGCGGTTGCCAAGGCATTGA<br>CGCTCGGATTGATCCGTTTGTGA | |
| *luxR* | CDS | ATGAAAAACATAAATGCCGACGACACA<br>TACAGAATAATTAATAAAATTAAAGCTT<br>GTAGAAGCAATAATGATATTAATCAATG<br>CTTATCTGATATGACTAAAATGGTACAT<br>TGTGAATATTATTTACTCGCGATCATTT<br>ATCCTCATTCTATGGTTAAATCTGATAT<br>TTCAATCCTAGATAATTACCCTAAAAAA<br>TGGAGGCAATATTATGATGACGCTAAT<br>TTAATAAAATATGATCCTATAGTAGATT<br>ATTCTAACTCCAATCATTCACCAATTAA<br>TTGGAATATATTTGAAAACAATGCTGTA<br>AATAAAAAATCTCCAAATGTAATTAAAG<br>AAGCGAAAACATCAGGTCTTATCACTG<br>GGTTTAGTTTCCCTATTCATACGGCTAA<br>CAATGGCTTCGGAATGCTTAGTTTTGC<br>ACATTCAGAAAAAGACAACTATATAGAT<br>AGTTTATTTTTACATGCGTGTATGAACA<br>TACCATTAATTGTTCCTTCTCTAGTTGA<br>TAATTATCGAAAAATAAATATAGCAAAT<br>AATAAATCAAACAACGATTTAACCAAAA<br>GAGAAAAAGAATGTTTAGCGTGGGCAT<br>GCGAAGGAAAAAGCTCTTGGGATATTT<br>CAAAAATATTAGGTTGCAGTGAGCGTA | 7 |

| | | | |
|---|---|---|---|
| | | CTGTCACTTTCCATTTAACCAATGCGC AAATGAAACTCAATACAACAAACCGCT GCCAAAGTATTTCTAAAGCAATTTTAAC AGGAGCAATTGATTGCCCATACTTTAA AAATTAA | |
| *rhlR* | CDS | ATGAGGAATGACGGAGGCTTTTTGCTG TGGTGGGACGGTTTGCGTAGCGAGAT GCAGCCGATCCACGACAGCCAGGGCG TGTTCGCCGTCCTGGAAAAGGAAGTG CGGCGCCTGGGCTTCGATTACTACGC CTATGGCGTGCGCCACACGATTCCCTT CACCCGGCCCAAGACCGAGGTCCATG GCACCTATCCCAAGGCCTGGCTGGAG CGATACCAGATGCAGAACTACGGGGC CGTGGATCCGGCGATCCTCAACGGCC TGCGCTCCTCGGAAATGGTGGTCTGG AGCGACAGCCTGTTCGACCAGAGCCG GATGCTCTGGAACGAGGCTCGCGATT GGGGCCTCTGTGTCGGCGCGACCTTG CCGATCCGCGCGCCGAACAATTTGCT CAGCGTGCTTTCCGTGGCGCGCGACC AGCAGAACATCTCCAGCTTCGAGCGC GAGGAAATCCGCCTGCGGCTGCGTTG CATGATCGAGTTGCTGACCCAGAAGCT GACCGACCTGGAGCATCCGATGCTGA TGTCCAACCCGGTCTGCCTGAGCCATC GCGAACGCGAGATCCTGCAATGGACC GCCGACGGCAAGAGTTCCGGGGAAAT CGCCATCATCCTGAGCATCTCCGAGAG CACGGTGAACTTCCACCACAAGAACAT CCAGAAGAAGTTCGACGCGCCGAACA AGACGCTGGCTGCCGCCTACGCCGCG GCGCTGGGCCTCATCTAA | 6 |
| *cinR* | CDS | ATGATTGAGAATACCTATAGCGAAAAG TTCGAGTCCGCGTTCGAACAGATCAAA GCGGCGGCCAACGTGGATGCCGCCAT CCGTATTCTCCAGGCGGAATATAACCT CGATTTCGTCACCTACCATCTCGCCCA GACAATCGCGAGCAAGATCGATTCGC CCTTCGTGCGCACCACCTATCCGGATG CCTGGGTTTCCCGTTACCTCCTCAACT GCTATGTGAAGGTCGATCCGATCATCA AGCAGGGCTTCGAACGCCAGCTGCCC TTCGACTGGAGCGAGGTCGAACCGAC GCCGGAGGCCTATGCCATGCTGGTCG ACGCCCAGAAACACGGCATCGATGAC | 8 |

| | | | |
|---|---|---|---|
| | | AATGGCTACTCCATCCCCGTCGCCGAC AAGGCGCAGCGCCGCGCCCTGCTGTC GCTGAATGCCCATATACCGGCCGACG AATGGACCGAGCTCGTGCGCCGCTGC CGCAATGAGTGGATCGAGATCGCCCA TCTGATCCACCGCAAGGCCGTATATGA GCTGCATGGCGAAAACGATCCGGTGC CGGCATTGTCGCCGCGCGAGATCGAG TGTCTGCACTGGACCGCCCTCGGCAA GGATTACAAGGATATTTCGGTCATCCT GGGCATATCAGAGCATACCACACGCG ATTACCTGAAAACCGCCCGCTTCAGGC TCGGCTGCACCACGATCTCGGCCGCC GCGTCGCGGGCTGTTCAATTGCGCAT CATCAATCCCTATAGGATCCGCATGAC GCGACGTAATTGGTAA | |
| *gfp* | CDS | ATGAGTAAAGGAGAAGAACTTTTCACT GGAGTTGTCCCAATTCTTGTTGAATTA GATGGTGATGTTAATGGGCACAAATTT TCTGTCAGTGGAGAGGGTGAAGGTGA TGCAACATACGGAAAACTTACCCTTAA ATTTATTTGCACTACTGGAAAACTACCT GTTCCATGGCCAACACTTGTCACTACT TTCGCGTATGGACTTCAATGCTTTGCG AGATACCCAGATCATATGAAACAGCAT GACTTTTTCAAGAGTGCCATGCCCGAA GGTTATGTACAGGAAAGAACTATATTTT TCAAAGATGACGGGAACTACAAGACAC GTGCTGAAGTCAAGTTTGAAGGTGATA CCCTTGTTAATAGAATCGAGTTAAAAG GTATTGATTTTAAAGAAGATGGAAACAT TCTTGGACACAAATTGGAATACAACTAT AACTCACACAATGTATACATCATGGCA GACAAACAAAAGAATGGAATCAAAGTT AACTTCAAAATTAGACACAACATTGAAG ATGGAAGCGTTCAACTAGCAGACCATT ATCAACAAAATACTCCAATTGGCGATG GCCCTGTCCTTTTACCAGACAACCATT ACCTGTCCTACCAATCTGCCCTTTCGA AAGATCCCAACGAAAAGAGAGATCACA TGGTCCTTCTTGAGTTTGTAACAGCTG CTGGGATTACACATGGCATGGATGAAC TATACAAATAA | 3 |
| *cinI* | CDS | ATGTTCGTTATCATTCAGGCACATGAG TATCAGAAATACGCTGCCGTACTCGAC CAGATGTTTCGTCTGCGCAAGAAGGTG | This study (*cinI* from *Rhizobium* |

| | | | |
|---|---|---|---|
| | | TTCGCCGATACGCTCTGCTGGGACGTT CCTGTCATCGGCCCTTACGAACGTGAC AGCTACGATTCGCTTGCTCCCGCCTAT CTCGTCTGGTGCAACGACAGCCGCAC CCGTCTTTATGGCGGCATGCGCCTGAT GCCGACGACCGGCCCGACCCTTCTCT ACGACGTGTTCCGCGAGACGTTCCCT GATGCCGCCGATCTTATCGCCCCCGG CATCTGGGAAGGCACGCGCATGTGCA TCGACGAGGAGGCGATCGCCAAGGAT TTCCCCGAGATCGACGCCGGCCGCGC CTTCTCCATGATGCTGCTCGCGCTTTG CGAATGCGCGCTCGATCACGGCATCC ACACGATGATCTCCAACTACGAGCCCT ACCTCAAGCGCGTCTACAAGCGCGCC GGCGCCGAGGTGGAAGAACTCGGCCG CGCAGACGGCTACGGCAAATATCCCG TCTGCTGCGGCGCCTTCGAAGTCTCG GACCGCGTGCTGCGCAAGATGCGCGC CGCCCTCGGCCTCACCCTACCCCTTTA TGTCAGGCACGTGCCGGCCCGCTCGG TCGTGACCCAATTCCTGGAGATGGCAG CATAA | *leguminosa-rum* with 2 nt mutated to remove two BbsI sites) |
| *luxI* | CDS | ATGACTATAATGATAAAAAAATCGGATT TTTTGGCAATTCCATCGGAGGAGTATA AAGGTATTCTAAGCCTTCGTTATCAAGT GTTTAAGCAAAGACTTGAGTGGGACTT AGTTGTAGAAAATAACCTTGAATCAGAT GAGTATGATAACTCAAATGCAGAATAT ATTTATGCTTGTGATGATACTGAAAATG TAAGTGGATGCTGGCGTTTATTACCTA CAACAGGTGATTATATGCTGAAAAGTG TTTTTCCTGAATTGCTTGGTCAACAGA GTGCTCCCAAAGATCCTAATATAGTCG AATTAAGTCGTTTTGCTGTAGGTAAAAA TAGCTCAAAGATAAATAACTCTGCTAGT GAAATTACAATGAAACTATTTGAAGCTA TATATAAACACGCTGTTAGTCAAGGTAT TACAGAATATGTAACAGTAACATCAACA GCAATAGAGCGATTTTTAAAGCGTATT AAAGTTCCTTGTCATCGTATTGGAGAC AAAGAAATTCATGTATTAGGTGACACTA AATCGGTTGTATTGTCTATGCCTATTAA TGAACAGTTTAAAAAAGCAGTCTTAAAT TAA | 10 |

| | | | |
|---|---|---|---|
| *rhlI* | CDS | ATGATTGAACTGTTAAGCGAGAGCTTA<br>GAGGGACTTAGCGCTGCGATGATTGC<br>GGAACTTGGGCGTTATCGCCACCAAGT<br>CTTTATTGAGAAACTCGGATGGGATGT<br>TGTTAGCACTAGCCGCGTGCGTGATCA<br>AGAGTTTGATCAATTTGATCACCCCCA<br>GACTCGTTATATTGTAGCGATGTCGCG<br>TCAAGGGATTTGTGGCTGTGCACGTCT<br>TTTACCTACTACTGATGCATATTTGCTG<br>AAAGATGTCTTTGCGTATTTATGTTCCG<br>AGACTCCACCCTCAGACCCCAGCGTTT<br>GGGAACTGAGTCGTTATGCGGCGTCC<br>GCTGCAGATGACCCCCAACTTGCTATG<br>AAAATCTTTTGGAGCTCATTGCAGTGT<br>GCGTGGTATCTCGGTGCATCTAGCGTT<br>GTTGCCGTTACGACGACAGCGATGGA<br>ACGTTACTTTGTGCGTAATGGTGTAATT<br>CTGCAACGTCTTGGTCCGCCACAAAAG<br>GTTAAAGGAGAAACCCTTGTTGCTATT<br>AGTTTTCCAGCATATCAAGAACGTGGT<br>TTGGAAATGCTCCTTCGTTATCATCCA<br>GAGTGGTTACAAGGTGTCCCATTGAGT<br>ATGGCTGTCTGA | This study<br><br>(*rhlI* from *P. aeruginosa* codon optimized for *E. coli* without BsaI, BbsI, or SapI cut sites) |
| L3S2P21 | Terminator | CTCGGTACCAAATTCCAGAAAAGAGGC<br>CTCCCGAAAGGGGGGCCTTTTTTCGTT<br>TTGGTCC | 9 |
| L3S2P15 | Terminator | CTCGGTACCAAATTCCAGAAAAGAGAC<br>GCTTTTAGAGCGTCTTTTTTCGTTTTGG<br>TCC | 9 |
| *amyE* front | Homology arm | ATGTTTGCAAAACGATTCAAAACCTCTT<br>TACTGCCGTTATTCGCTGGATTTTTATT<br>GCTGTTTCATTTGGTTCTGGCAGGACC<br>GGCGGCTGCGAGTGCTGAAACGGCGA<br>ACAAATCGAATGAGCTTACAGCACCGT<br>CGATCAAAAGCGGAACCATTCTTCATG<br>CATGGAATTGGTCGTTCAATACGTTAA<br>AACACAATATGAAGGATATTCATGATG<br>CAGGATATACAGCCATTCAGACATCTC<br>CGATTAACCAAGTAAAGGAAGGGAATC<br>AAGGAGATAAAAGCATGTCGAACTGGT<br>ACTGGCTGTATCAGCCGACATCGTATC<br>AAATTGGCAACCGTTACTTAGGTACTG<br>AACAAGAATTTAAAGAAATGTGTGCAG<br>CCGCTGAAGAATATGGCATAAAGGTCA<br>TTGTTGACGCGGTCATCAATCATACCA<br>CCAGTGATTATGCCGCGATTTCCAATG | 3 |

| | | | |
|---|---|---|---|
| | | AGGTTAAGAGTATTCCAAACTGGACAC<br>ATGGAAACACACAAATTAAAAACTGGT<br>CTGATCGATGGGATGTCACGCAGAATT | |
| *amyE* back | Homology arm | ATCCGTTTAGGCTGGGCGGTGATAGCT<br>TCTCGTTCAGGCAGTACGCCTCTTTTC<br>TTTTCCAGACCTGAGGGAGGCGGAAAT<br>GGTGTGAGGTTCCCGGGGAAAAGCCA<br>AATAGGCGATCGCGGGAGTGCTTTATT<br>TGAAGATCAGGCTATCACTGCGGTCAA<br>TAGATTTCACAATGTGATGGCTGGACA<br>GCCTGAGGAACTCTCGAACCCGAATG<br>GAAACAACCAGATATTTATGAATCAGC<br>GCGGCTCACATGGCGTTGTGCTGGCA<br>AATGCAGGTTCATCCTCTGTCTCTATC<br>AATACGGCAACAAAATTGCCTGATGGC<br>AGGTATGACAATAAAGCTGGAGCGGG<br>TTCATTTCAAGTGAACGATGGTAAACT<br>GACAGGCACGATCAATGCCAGGTCTG<br>TAGCTGTGCTTTATCCTGATGATATTGC<br>AAAAGCGCCTCATGTTTTCCTTGAGAA<br>TTACAAAACAGGTGTAACACATTCTTTC<br>AATGATCAACTGACGATTACCTTGCGT<br>GCAGATGCGAATACAACAAAAGCCGTT<br>TATCAAATCAATAATGGACCAGAGACG<br>GCGTTTAAGGATGGAGATCAATTCACA<br>ATCGGAAAAGGAGATCCATTTGGCAAA<br>ACATACACCATCATGTTAAAAGGAACG<br>AACAGTGATGGTGTAACGAGGACCGA<br>GAAATACAGTTTTGTTAAAAGAGATCCA<br>GCGTCGGCCAAAACCATCGGCTATCAA<br>AATCCGAATCATTGGAGCCAGGTAAAT<br>GCTTATATCTATAAACATGATGGGAGC<br>CGAGTAATTGAATTGACCGGATCTTGG<br>CCTGGAAAACCAATGACTAAAAATGCA<br>GACGGAATTTACACGCTGACGCTGCCT<br>GCGGACACGGATACAACCAACGCAAA<br>AGTGATTTTTAATAATGGCAGCGCCCA<br>AGTGCCCGGTCAGAATCAGCCTGGCT<br>TTGATTACGTGCTAAATGGTTTATATAA<br>TGACTCGGGCTTAAGCGGTTCTCTTCC<br>CCATTGA | 3 |

Data in all supplementary tables is provided in the Supplementary Data file.

**Supplementary Table 4. DNA oligos used in this work**

| Oligo name | Sequence | Use |
|---|---|---|
| BS403 | AAGAAGACAACTCCAGCGACCTCATTGG | Forward primer to inverse PCR to remove $P_{FtsH}$ and add $P_{Bla}$ or $P_{Veg}$ in a subsequent DNA assembly |
| BS404 | AAGAAGACAACTGAAGTCATAAGTCTGGGCTAAG | Reverse primer to inverse PCR to remove $P_{FtsH}$ and add $P_{Bla}$ or $P_{Veg}$ in a subsequent DNA assembly |
| PBla_AS-O-Fw | GGAGTTGACACCGATATTACAATTGTAATATTATTGATTT | Forward oligo to make annealed DNA for $P_{Bla}$ |
| PBla_AS-O-Rv | TCAGAAATCAATAATATTACAATTGTAATATCGGTGTCAA | Reverse oligo to make annealed DNA for $P_{Bla}$ |
| Pveg_AS-O-Fw | GGAGTTGATATAATTTAAATTTTATTTGACAAAAATGGGCTCGTGTTGTACAATAAATGT | Forward oligo to make annealed DNA for $P_{Veg}$ |
| Pveg_AS-O-Rv | TCAGACATTTATTGTACAACACGAGCCCATTTTTGTCAAATAAAATTTAAATTATATCAA | Reverse oligo to make annealed DNA for $P_{Veg}$ |
| BS303-Fw | AAGAAGACAACTAACAAACTCTTGGCTTGGAG | Forward primer to create backbone to insert HSL sensor constructs in pBS9000 vector |
| BS303-Rv | AAGAAGACAAGGAGGAGTTAACATTACCTGGACC | Forward primer to create backbone to insert HSL sensor constructs in pBS9000 vector |
